# Engineered *Pseudomonas putida* reconfigures metabolic fluxes to support energy demands during muconate bioproduction from lignin-related aromatics

**DOI:** 10.64898/2026.07.14.738580

**Authors:** Rebecca A. Wilkes, Patrick F. Suthers, Andrew J. Borchert, Melanie M. Callaghan, Eashant Thusoo, Richard J. Giannone, Dana L. Carper, John I. Hendry, Alexander F. Benson, Michael A. Gapuz, Ashley N. Merrill, Kelsey J. Ramirez, Davinia Salvachúa, Robert L. Hettich, Costas D. Maranas, Daniel Amador-Noguez, Gregg T. Beckham, Allison Z. Werner

## Abstract

Muconic acid is a versatile platform chemical that can be biologically produced from lignocellulosic substrates, including from lignin-related aromatic compounds. *Pseudomonas putida* has been previously engineered to convert lignin-related aromatic compounds to muconate at quantitative molar yields. This high atom efficiency requires a supplemental carbon and energy source to support bacterial growth, and central carbon metabolic efficiency and its interaction with aromatic catabolism are underexplored. Here, we applied proteomics, metabolomics, and ^13^C-fluxomics to quantitatively compare central carbon and energy metabolism in wild-type *P. putida* KT2440 and a muconate-producing strain, *P. putida* CJ781. During cultivation on glucose and 4-hydroxybenzoate, CJ781 showed increased glucose uptake, reconfigured central fluxes, and increased extracellular leakage of aliphatic acids relative to wild type. These altered fluxes supported a 3-fold higher ATP pool, in excess of demand. Pyruvate and acetate secretion in CJ781 was mitigated by debottlenecking TCA-cycle entry via citrate synthase overexpression. Furthermore, tuned expression of the catechol dioxygenase and protocatechuate decarboxylase enabled the production of 36.3 g L^-1^ muconate at 1.1 g L^-1^ h^-1^. Overall, this work reveals how *P. putida* redirects carbon and energy fluxes to support aromatic bioconversion for improved bioproduction from renewable feedstocks.

## INTRODUCTION

Multiple bioproducts are accessible from lignin-related aromatic compounds at atom-efficient yields,^1–5^ including *cis, cis*-muconic acid (hereafter muconate),^6–10^ β-ketoadipic acid,^11–15^ and 2-pyrone-4,6-dicarboxylic acid.^16,17^ Muconate has gained particular traction as it can be used as a direct replacement to petrochemicals such as adipic acid^18–20^ or converted to performance-advantaged bioproducts,^21–26^ making it a bioprivileged molecule.^27^ Lignin-related hydroxybenzoates, such as 4-hydroxybenzoate (4HB) and vanillate, can be rerouted to muconate via AroY-mediated protocatechuate (PCA) decarboxylation to catechol^9,28^ followed by ring-opening to muconate via the catechol 1,2-dioxygenase CatA (**Fig. 1**). Deletion of *pcaHG* and *catBC* prevents aromatic carbon conversion to succinate and acetyl-CoA and subsequently into biomass and energy production.^28^

**Figure 1.**
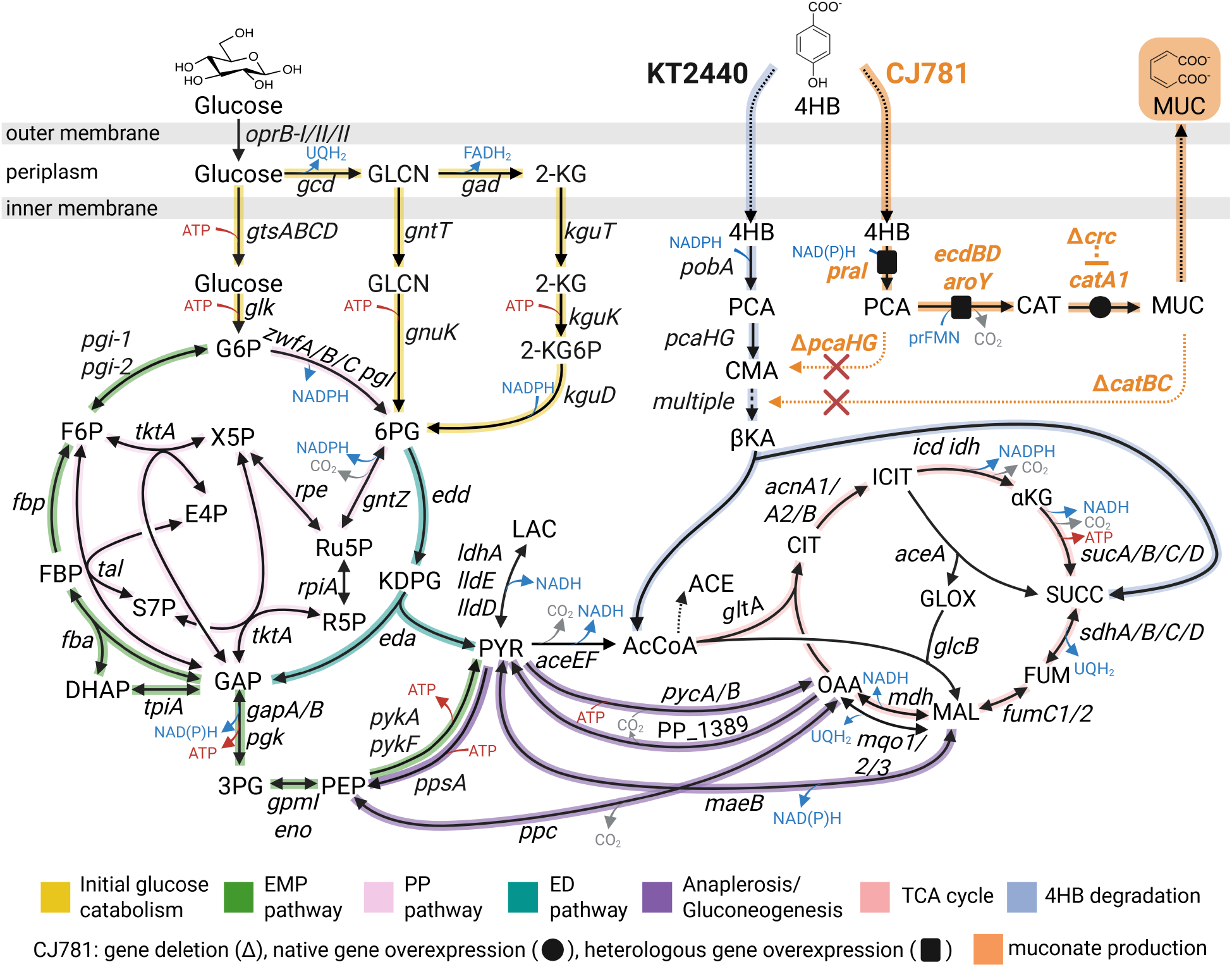
Pathway map for glucose metabolism and either 4HB catabolism to central carbon pathways (KT2440) or conversion to muconate (CJ781). Metabolite abbreviations: 4HB, 4-hydroxybenzoate; PCA, protocatechuate; CMA, β-carboxy-*cis,cis*-muconate; AcCoA, acetyl-CoA; CA, catechol; MA, *cis,cis*-muconate; GLCN, gluconate; 2-KG, 2-ketogluconate; P, phosphate; 2-KG6P, 2-ketogluconate-6-P; G6P, glucose-6-P; 6PG, 6-phosphogluconate; KDPG, 2-keto-3-deoxy-6-phosphogluconate; GAP, glyceraldehyde-3-P; FBP, fructose-1,6-bisP; F6P, fructose-6-P; S7P, sedoheptulose-7-P; R5P, ribose-5-P; Ru5P, ribulose-5-P; X5P, xylulose-5-P; E4P, erythrose 4-P; 3PG, 3-phosphoglycerate; PEP, phosphoenolpyruvate; PYR, pyruvate; LAC, lactate; ACE, acetate; CIT, citrate; ICIT isocitrate; αKG, alpha-ketoglutarate; SUCC, succinate; FUM, fumarate; MAL, malate; OAA, oxaloacetate; GLOX, glyoxylate.

*Pseudomonas putida* KT2440 (hereafter KT2440) has emerged as a platform strain for aromatic bioconversion, largely owing to its established genetic engineering tools,^29–31^, catabolic versatility,^8,14,32–35^ stress tolerance,^36–39^ and capacity to produce a variety of products at high titers.^3,29^ Over the past decade, our group has improved muconate production from hydroxybenzoates in KT2440 by deleting the *crc* catabolite repressor, overexpressing the vanillate monooxygenase encoded by *vanAB*, overexpressing *ecdBD*, which is associated with the production of the AroY cofactor prenylated flavin mononucleotide (prFMN),^40^ and replacing the native 4HB hydroxylase that requires NADPH (*pobA*) with a heterologous homolog that accepts either NADPH or NADH (*praI*).^10,28,41,42^ The culmination of these modifications, resulting in strain CJ781 (**Fig. 1, Table 1**), enabled muconate production from *p-*coumarate with supplemental glucose at a productivity of 0.5 g L^-1^ h^-1^, a titer of 40 g L^-1^, and a molar yield of 100%.^10^ However, productivity greater than 1 g L^-1^ h^-1^ remains a significant barrier to economical strain performance,^15,43^ and thus additional efforts are required to improve the efficiency of muconate production further.

**Table 1.**
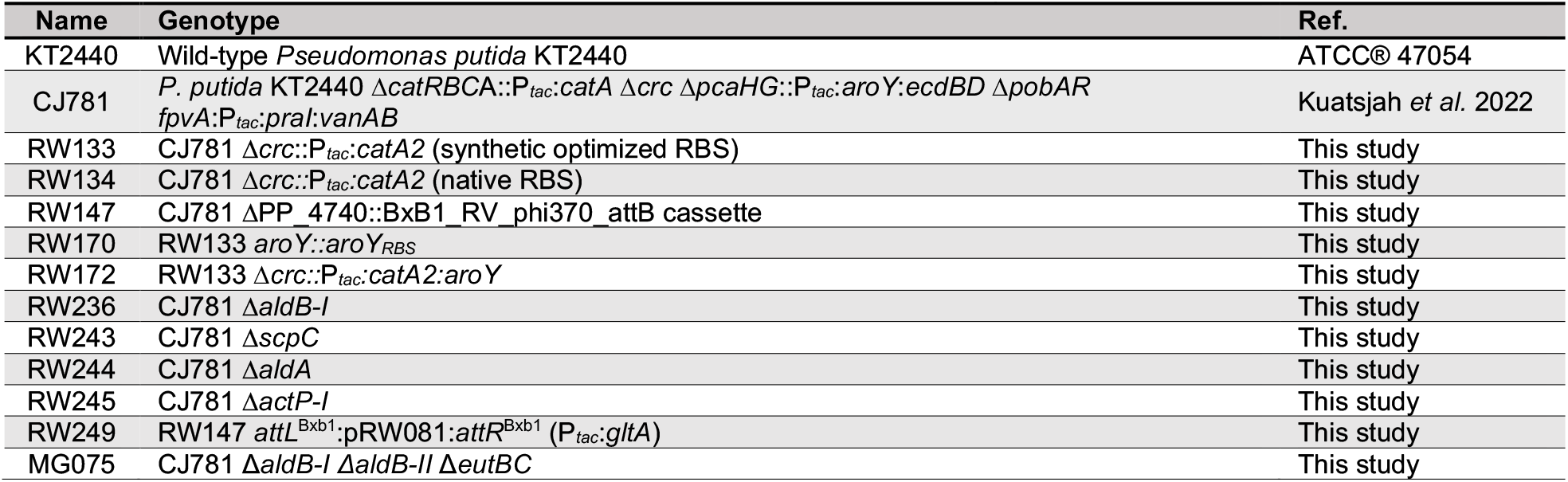
*P. putida* strains used in this study. Abbreviation: RBS, ribosome binding site.

A supplemental carbon and energy source is required to support cellular growth and maintenance in CJ781, as aromatic-derived carbons are diverted to product instead of being incorporated into biomass (**Fig. 1**).^10,15,44–47^ Glucose is a commonly used carbon source, and its metabolism in KT2440 has been well-studied and known to undergo periplasmic oxidation reactions to rapidly produce reducing equivalents without ATP expenditure.^48–50^ However, the impact of product formation on central carbon and energy metabolism is understudied yet critical,^45^ as engineered biosynthesis pathways often compete with native physiology.^51,52^ During muconate production from catechol, a KT2440-derived strain decreased periplasmic glucose oxidation in favor of the glucose-phosphorylation route, increasing NADPH supply.^45^ However, the opposite trend of elevated periplasmic glucose oxidation and high ATP excess over biomass demand was observed from heterologous protein expression in KT2440.^53^ Thus, the impact of aromatic substrate conversion to muconate on glucose metabolism and cellular energetics warrants further investigation.

In this work, we combined proteomics, metabolomics, and ^13^C-fluxomics to understand the impact of heterologous pathway engineering and muconate production on glucose metabolism by characterizing KT2440 and CJ781 fed with glucose only or glucose plus 4HB. During muconate production, CJ781 changed carbon and energy fluxes, accumulated intracellular metabolites, and increased aliphatic acid secretions compared to KT2440. These data informed our engineering strategies to both decrease pyruvate and acetate secretions and resolve aromatic catabolic bottlenecks, ultimately delivering strains that both enabled higher 4HB feeding rates and produced muconate at greater than 2-fold higher rates than CJ781.

## RESULTS

### Integrated ^13^C-fluxomics, metabolomics, and proteomics analysis of wild type and muconate production strain

We first sought to understand metabolic rewiring and energetics in the engineered strain CJ781 (**Fig. 1, Table 1**), under conditions with and without muconate production, as compared to wild-type KT2440. To mitigate toxicity responses from aromatic acids and focus on changes specific to muconate production, we chose to use a low concentration (5 mM) of the least toxic relevant aromatic compound, 4HB.^42^ Accordingly, we cultivated KT2440 and CJ781 in M9 minimal media supplemented with 20 mM glucose alone or 20 mM glucose plus 5 mM 4HB (**Fig. 2a**). To allow cells to acclimate to the test conditions used, seed cultures were grown for 10-12 h in minimal media containing the experimental substrates prior to inoculating experimental cultures for measurement.

**Figure 2.**
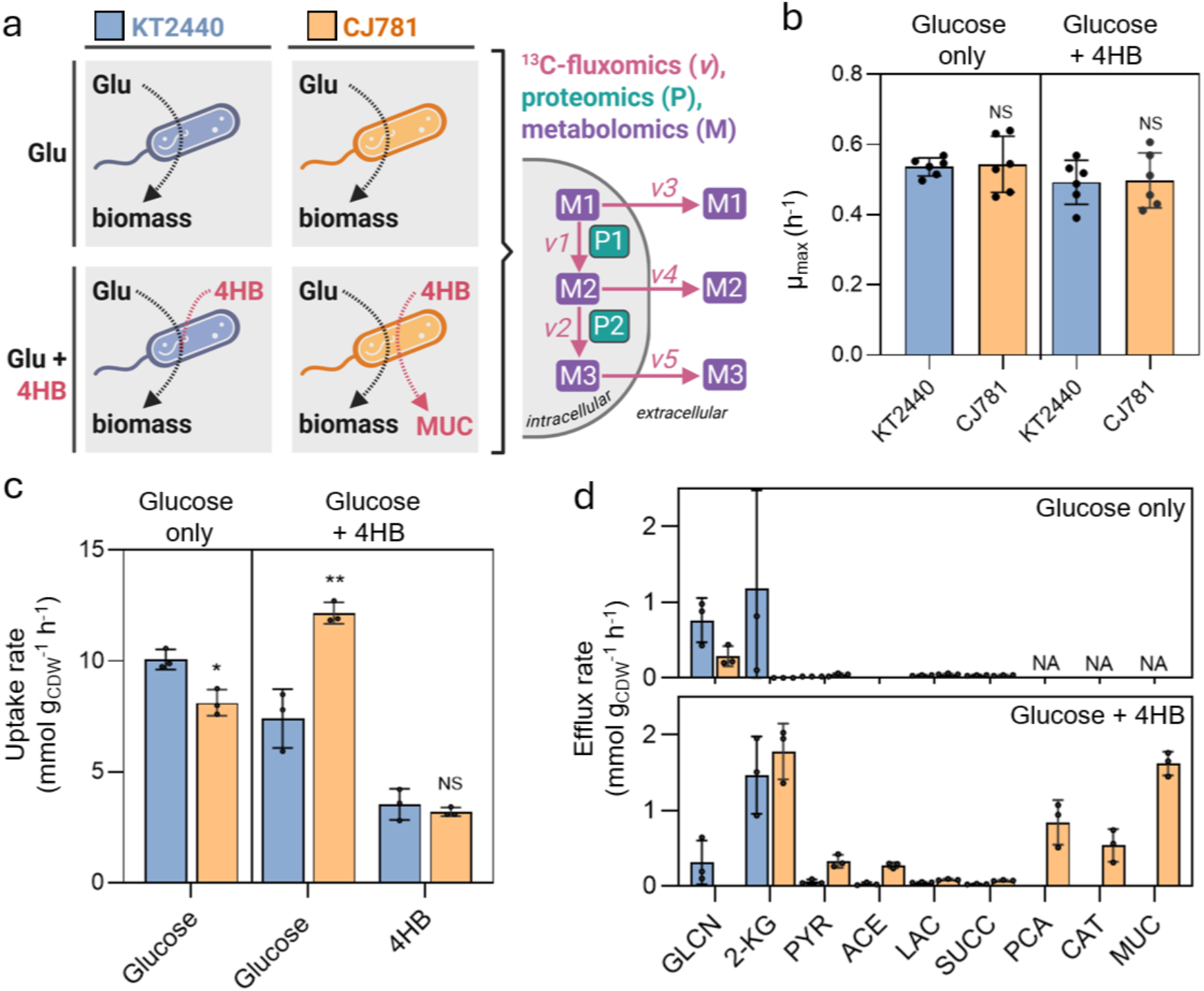
Growth and carbon utilization of wild type (KT2440) and a muconate production strain (CJ781) on glucose alone or glucose plus lignin-related aromatic 4HB. (**a**) Schematic of the approach wherein KT2440 (blue grey) and CJ781 (orange) were cultivated on either glucose alone or glucose plus 4HB. The partitioning of the aromatic catabolic pathway to muconate from glucose metabolism is highlighted for CJ781. Each strain and condition were analyzed by proteomics, metabolomics, and ^13^C-fluxomics. (**b**) Growth rates for KT2440 and CJ781 on glucose only or glucose plus 4HB. The data represent the mean ± the standard deviation determined from six independent biological replicates. (**c**) Uptake rates of glucose and 4HB calculated during exponential growth. (**d**) Metabolite secretion rates of compounds measured in the extracellular milieu. For **c** and **d**, the data represent the mean ± the standard deviation determined from three independent biological replicates. Statistically significant differences (p < 0.05) were determined with an unpaired two-tailed *t*-test between CJ781 and KT2440. Rates are reported in **Table S1**. All data used to calculate rates for **b, c**, and **d** are available in **Excel file 1**. Metabolite abbreviations are listed in **Fig. 1**.

Despite the inability of CJ781 to convert 4HB to biomass, the growth rate was comparable between CJ781 and KT2440 in both the presence and absence of 4HB (**Fig. 2b; Table S1**). In the presence of 4HB, CJ781 exhibited a ∼2 h longer lag than KT2440 but the lag was similar for both strains on glucose alone (**Fig. S1**). Additionally, the mass (g) of protein per mass (g) cell dry weight (g_CDW_) was comparable between all the strains and conditions, indicating that overexpression of the five protein-encoding genes in CJ781 did not alter the overall biomass protein allocation (**Fig. S2**). In contrast, the g of biomass per g of glucose consumed (Y_x/glu_) differed for CJ781 relative to KT2440. On glucose alone, CJ781 had a 25% greater Y_x/glu_ (*p* = 0.014; unpaired two-tailed *t*-test) than KT2440, whereas, in the presence of 4HB, CJ781 had 40% lower Y_x/glu_ (*p* = 0.025; unpaired two-tailed *t*-test) than KT2440 (**Table S1**), signifying differences in glucose conversion to biomass during muconate production.

To investigate these differences in glucose metabolism, we performed ^13^C-metabolic flux analysis (fluxomics), metabolomics, and proteomics on both strains under both conditions (**Fig. 2a, Table 1**). For all three analyses, cells were harvested during exponential growth, when glucose and 4HB were depleted simultaneously (**Fig. S3**). Principal component analysis of the proteomes from the two strains and growth conditions demonstrated that there was close agreement between biological replicates, with data variation driven by genotype and substrate composition (**Fig. S4**). For high-resolution ^13^C-fluxomics, parallel tracer experiments were conducted with 100% [1-^13^C], 100% [6-^13^C], and 50:50% [U-^13^C_6_] and unlabeled glucose for both strains and media compositions. The labeling patterns of central carbon metabolites and amino acids were analyzed for up to 321 total mass isotopomers during the high-resolution flux analysis of each strain and substrate composition pairing (**Excel file 1**). Biomass and extracellular metabolites were measured every 1-3 h throughout the time-course to calculate the uptake and efflux rates used to constrain the model (**Fig. 2b-d; Excel file 1**). For all conditions, the model estimation for metabolic flux analysis reached statistically significant model fits and low sum of squared weighted residuals to the effluxes and ^13^C-labeling data (**Table S2; Fig. S5**).

### Rates for glucose uptake and oxidation to 2-ketogluconate increase during muconate production

We first compared glucose and 4HB uptake rates. During cultivation on glucose plus 4HB, CJ781 had a similar 4HB uptake rate but the glucose uptake rate increased by 64% compared to KT2440 (**Fig. 2c**). OprB-I, a permease that transports glucose into the periplasm, was concomitantly increased in abundance by 2.2-fold (**Fig. 3b**). Interestingly, the glucose uptake rate alone, in carbon equivalents, for CJ781 (72.9 ± 12.5 mM carbon g _CDW_ ^-1^ h^-1^) resembled the cumulative carbon uptake rate of glucose and 4HB (69.2 ± 11.3 mM carbon g_CDW_ ^-1^ h^-1^) for KT2440 (**Fig. S6**). Conversely, when fed glucose only, CJ781 had a 20% reduction in the glucose uptake rate compared to KT2440 (**Fig. 2c**). Thus, the increased uptake rate for CJ781 observed during co-feeding glucose and 4HB cannot be attributed solely to genotype but rather reflects an interplay between environmental response and genetic background.

**Figure 3.**
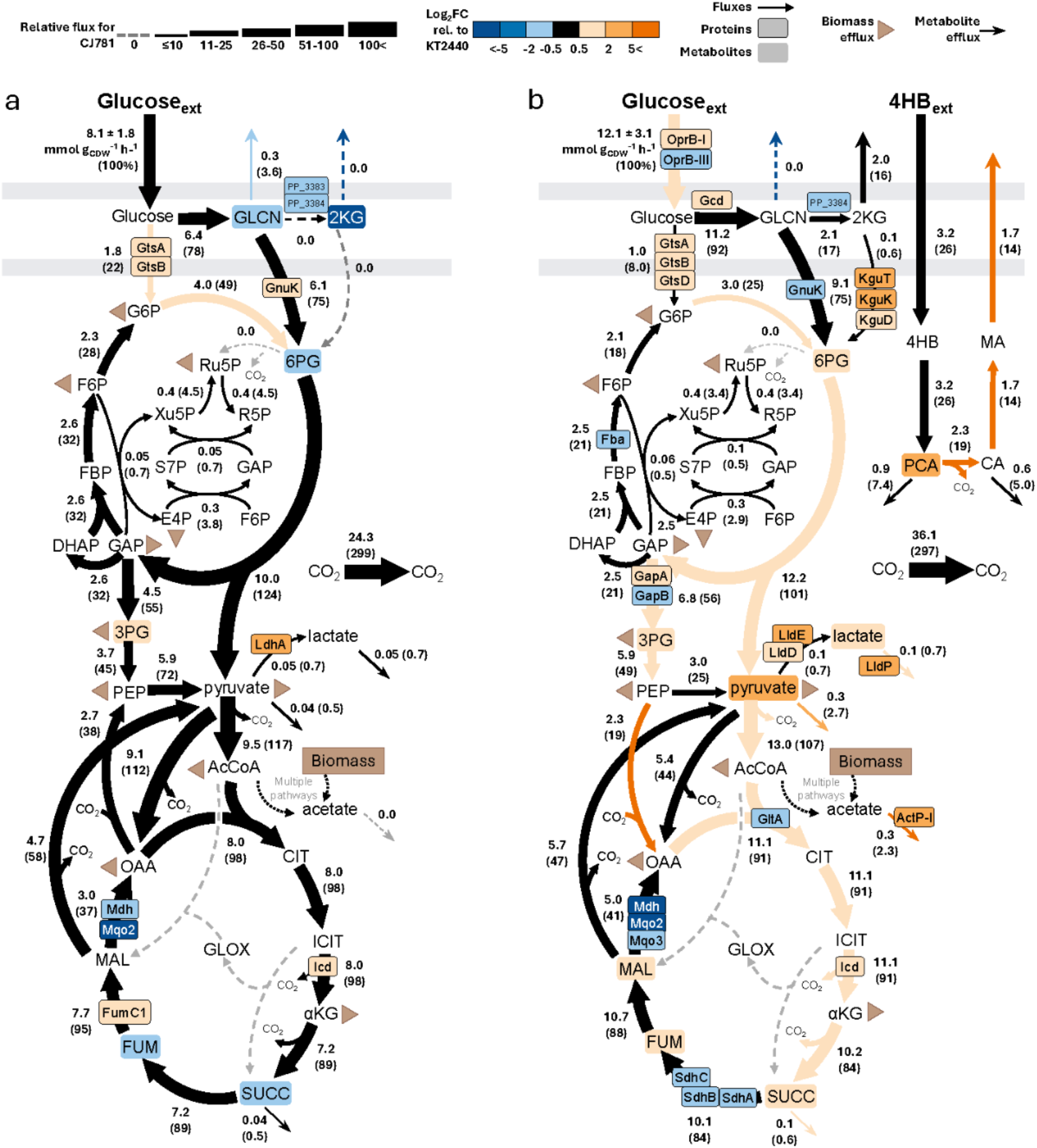
Integrated proteomics, metabolomics, and metabolic flux modeling. ^13^C-metabolic fluxes for CJ781 fed (**a**) glucose only or (**b**) glucose plus 4HB. For both **a** and **b**, absolute fluxes for CJ781, in mmol g_CDW_ ^-1^ h^-1^, are denoted along with the relative fluxes as a percentage of glucose uptake, shown in parentheses and depicted by arrow width. Differentially abundant proteins and metabolites (FDR < 0.05 for proteomics; p < 0.05 for metabolomics) with log_2_ fold-change (FC) relative to KT2440 that were either greater than 0.5 or less than -0.5 are highlighted in the metabolic map. Fluxes that had 95% CI intervals that did not overlap are also highlighted. Metabolic fluxes were optimized using three parallel labeling schemes and a total of nine independent biological replicates. Proteins and metabolites were measured from three independent biological replicates. Data for fluxes, metabolites, and proteins are provided in **Excel file 1**. Abbreviations are listed in **Fig. 1**.

We next considered changes in periplasmic glucose oxidation and secretion of gluconate and 2-ketogluconate (2-KG), as differences in glucose metabolism via these oxidation and phosphorylation pathways can rapidly generate different energy and redox stoichiometry (**Fig. 1**). Notably, there was low precision in gluconate and 2-KG fluxes as they were constrained solely by gluconate and 2-KG secretions (**Excel file 1**), so a combination of the protein, metabolite, and flux data was used to examine these pathways (**Fig. 2d, Fig. 3**).

On glucose alone, CJ781 exhibited a low gluconate secretion rate (3.0 ± 1.6% of glucose uptake rate) and no measurable 2-KG secretion (**Fig. 2d**). Increased siphoning of glucose into the cytosol through glucose phosphorylation was corroborated for CJ781 by the elevated transporters (GtsA, GtsB; >1.8-fold increase) and flux from glucose to glucose-6-phosphate (G6P) (3.8-fold increase) compared to KT2440 (**Fig. 3a**). In addition, GnuK increased 1.5-fold, which would support flux through the gluconate phosphorylation pathway, and there were also lower periplasmic gluconate dehydrogenases PP_3383 and PP_2284 (2.6- and 3.4-fold decrease), depleted intracellular pools of gluconate and 2-KG (3.9- and >1,000-fold decrease), and lower gluconate secretion (2.7-fold decrease) (**Fig. 3a**). Thus, CJ781 on glucose alone had less flux through periplasmic oxidation reactions to 2-KG, and elevated uptake through glucose phosphorylation, potentially offsetting the lower glucose uptake rate compared to KT2440 (**Fig. 2c**).

In contrast, some of the changes for CJ781 relative to KT2440 were reversed in the presence of 4HB. During cultivation on glucose and 4HB, CJ781 secreted 2-KG at 14.6 ± 3.0% of the glucose uptake rate and had no measurable gluconate secretion rate (**Fig. 2d**). Although GtsA and GtsB remained elevated similar to when fed glucose only, the glucose dehydrogenase (Gcd) that initiates periplasmic oxidation to gluconate increased 1.7-fold for CJ781 relative to KT2440 on glucose and 4HB (**Fig. 3a-b**). Elevated protein abundances were also observed for the 2-KG importer (KguT, 25-fold increase) and enzymes converting 2-KG to 6-phosphogluconate (6PG) (KguK and KguD; 6.3- and 3.8-fold increases, respectively), as well as a 2.2-fold increased 6PG pool (**Fig. 3**). KguT, KguK, and KguD were also elevated up to 3.4-fold for CJ781 in the presence of 4HB relative to glucose alone (**Fig. S7**). These protein-level and metabolite-level changes indicated that CJ781 exhibited increased flux to 2-KG during muconate production, potentially supporting rapid energy production via the periplasmic oxidation reactions that produce UQH_2_ and FADH_2_.

### Muconate production is accompanied by central carbon metabolite accumulation and secretion

KT2440 processes glucose through portions of the Entner-Doudoroff (ED), Embden-Meyerhof-Parnas (EMP), and oxidative pentose phosphate (PP) pathways in a cycle called the cyclic EDEMP pathway that fuels NADPH regeneration (**Fig. 1**).^29,48,54^ Pyruvate produced from the EMP and ED pathway acts as a link between these pathways and the TCA cycle, which is central to generating reduced cofactors and energy in the cell (**Fig. 1**). Curiously, we observed both intracellular accumulation and secretion of central carbon aliphatic acids in CJ781 (**Fig. 2d, Fig. 3, Fig. S8**). During cultivation on glucose plus 4HB, intracellular metabolite pools for 6PG, 3-phosphoglycerate (3PG), lactate, succinate, fumarate, malate, and pyruvate were elevated by at least 1.8-fold in CJ781 compared to KT2440 (**Fig. 3b**). In particular, the intracellular pool of pyruvate was 6.1-fold greater, and the efflux rate was 5.7-fold higher for CJ781 relative to KT2440, highlighting pyruvate as a substantial carbon surplus and a potential metabolic bottleneck in glucose metabolism during muconate production. The citrate synthase GltA was depleted by 1.9-fold in CJ781 relative to KT2440, potentially decreasing flux into the TCA cycle and exacerbating pyruvate accumulation.

Remarkably, 40 transport proteins, including organic acid exporters, were significantly elevated in CJ781 relative to KT2440 in the presence of 4HB (**Fig. 3, Fig. S9**). For example, we observed a 11.6-fold increase in abundance for the lactate exporter LldP and a 5.2-fold increase in abundance for the acetate transporter ActP-I. Additionally, the lactate dehydrogenases LldE and LldD were increased in abundance by 4.1- and 3.5-fold, respectively, facilitating the 1.7-fold increase in lactate flux (**Fig. 3**). However, after accounting for the carbon uptake apportioned to metabolite secretions and biomass demand, CJ781 had equivalent carbon flux remaining for CO_2_ as KT2440 fed glucose and 4HB (**Fig. S10**).

Conversely, organic acid secretion rates remained <1% of the glucose uptake rate during growth on glucose alone for both CJ781 and KT2440 (**Fig. 2d, Fig. 3**). Similarly, only 10 transport proteins were increased in CJ781 relative to KT2440 in the absence of 4HB (**Fig. S9**). Overall, these data indicate that substrate composition supporting muconate production, rather than the CJ781 genotype alone, underlies the observed accumulation and secretion of central carbon metabolites, indicative of changes to protein allocation and carbon fluxes to maintain cell growth and energy requirements.

### CJ781 increased fluxes through ED, gluconeogenic, and TCA cycle pathways in presence of 4HB

Across both strains and substrate compositions, the net fluxes, in mmol g_CDW_^-1^ h^-1^, were consistent through the non-oxidative PP pathway, the gluconeogenic EMP portion of the EDEMP cycle, and the pyruvate shunt [malate to pyruvate to oxaloacetate (OAA)], implying an optimized flux through these pathways to maintain biomass precursors and reduced cofactors (**Fig. 3; Excel File 1**). The genetic background for CJ781 relative to KT2440 had a few consistent differences independent of substrate compositions, including elevations in isocitrate dehydrogenase abundance (Icd, >1.5-fold), increased flux from glucose-6-phosphate (G6P) to 6-phosphogluconate (6PG) (∼1.5-fold), and depletion of malate dehydrogenase (Mdh) and malate-quinone oxidoreductase (Mqo2, >3-fold) (**Fig. 3**; **Excel file 1**), which would all support NADPH regeneration. Additional differences across central carbon metabolism were identified for CJ781 relative to KT2440 in the presence of 4HB (**Fig. 3b**).

The net fluxes through the ED pathway and oxidative side of the TCA cycle increased by 1.5-fold in CJ781 relative to KT2440 during cultivation on glucose plus 4HB (**Fig. 3b**). However, this higher relative flux was not observed on the reductive side of the TCA cycle. KT2440 assimilates 4HB-derived carbon into succinate and acetyl-CoA, which stimulated, relative to the glucose-only condition, the elevation of SdhABC (succinate dehydrogenase subunits) by >2-fold, FumC1 (fumarase) by 2.1-fold, and Mdh (malate dehydrogenase) by 7.3-fold (**Fig. S7**), supporting the 1.2-fold higher flux through the reductive side versus oxidative side of the TCA cycle (**Excel file 1**). In contrast, CJ781 does not assimilate 4HB-derived carbon, and SdhABC was depleted by >1.6-fold and Mdh was depleted 12.2-fold relative to KT2440 (**Fig. 3b**). Together, these data indicate that CJ781 rewires metabolism towards energy generation via elevated ED and oxidative TCA cycle flux during muconate production.

Phosphoenolpyruvate (PEP) carboxylase flux was reversed for CJ781 compared to KT2440 during muconate production, diverting carbon from PEP to OAA in the TCA cycle instead of toward pyruvate (**Fig. 3b**). This rewiring could act as a cellular response to the accumulation of pyruvate (**Fig. 3b, Fig. S8**). The PEP precursor, 3PG, was similarly elevated (>2.5-fold), in accordance with increased abundance of the glycolytic glyceraldehyde-3-phosphate dehydrogenase GapA (1.9-fold increase) and decreased abundance of the gluconeogenic isoform GapB (1.4-fold decrease) (**Fig. 3b**). Thus, CJ781 exhibited a drive for glycolytic and cataplerotic flux over gluconeogenic flux in the presence of 4HB.

### CJ781 generates an apparent excess NADPH and ATP, especially during muconate production

The impact of remodeled fluxes on cofactor and energy balances is key to understanding the availability of NADPH and ATP required for growth and stress responses. To obtain a quantitative understanding of energy metabolism, we balanced the supply and demand for NADPH, NADH, UQH_2_, and ATP from the ^13^C-fluxomics data (**Excel file 1, Fig. 4a-b**). NADPH supply was in excess of anabolic demand in all conditions but was highest in CJ781 during muconate production (19.8 ± 3.1 mmol NADPH g_CDW_ ^-1^ h^-1^). The ∼70% increase in apparent excess of NADPH in CJ781 during cultivation on glucose plus 4HB was primarily attributed to a ∼45% increase in flux through isocitrate dehydrogenase in the TCA cycle. The excess of NADPH quantified for each strain was assumed to undergo transhydrogenase conversion to NADH and then oxidative phosphorylation to produce ATP, adding to model-predicted generation of ATP. For ATP generation, CJ781 had a 1.6-fold higher ATP supply (137.3 ± 4.9 mmol ATP g_CDW_ ^-1^ h^-1^) and 2.5-fold increase in apparent ATP excess compared to KT2440 when fed glucose and 4HB (**Fig. 4b**). Both NADPH and ATP demand were consistent across all conditions, reflecting the similar growth rates across strains and media (**Fig. 2b, Fig. 4**).

**Figure 4.**
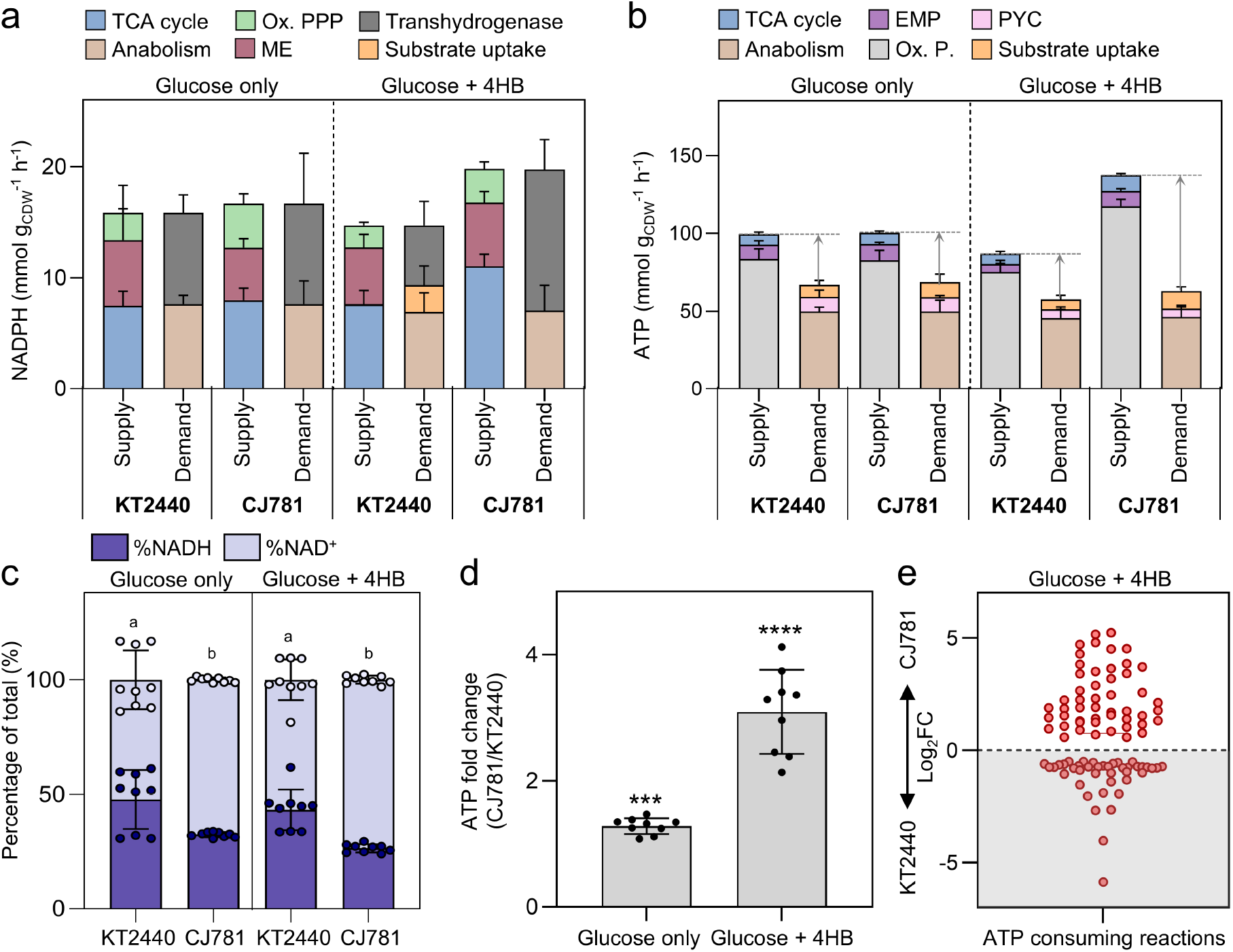
Quantitative analysis of redox cofactors and energy metabolism. (**a**) NADPH and (**b**) ATP supply and demand rates (mmol g_CDW_^-1^h^-1^) calculated from the obtained fluxes (**Fig. 3, Excel file 1)**. For **a** and **b**, error bars represent the cumulative standard deviation of the best-fit solution modeled across nine sets of ^13^C-labeling data. (**c**) Experimentally measured NAD^+^ and NADH pools as a percentage of the total NAD(H) pool during exponential growth. Error bars represent the standard deviation of the mean across three independent biological replicates and three technical replicates. A change in letter denotes a significant difference (p<0.05) determined from one-way ANOVA followed by Tukey HSD *post hoc* tests. (**d**) Experimentally measured fold change in ATP for CJ781 relative to KT2440 when fed glucose alone or glucose plus 4HB. Error bars represent the standard deviation of the mean across three independent biological replicates and three technical replicates. (**e**) Log_2_ fold-changes (FC) for ATP-consuming proteins measured during feeding glucose plus 4HB for CJ781 relative KT2440. Data were from three independent biological replicates. Abbreviations: Ox. PPP, oxidative pentose phosphate pathway; ME, malic enzyme; EMP, Embden-Meyerhof-Parnas pathway; PYC, pyruvate carboxylase; Ox. P, oxidative phosphorylation.

To experimentally confirm the high ATP supply in CJ781, we measured the NAD^+^/NADH and ATP/ADP ratios, and the total ATP pool (**Fig. 4c-d, Fig. S11**). KT2440 maintained an equivalent pool of NAD^+^ and NADH across both growth conditions, whereas CJ781 had higher NAD^+^ on both glucose alone and glucose plus 4HB (67.6 ± 1.1 % and 73.6 ± 1.8% of the total NAD(H) pool, respectively) (**Fig. 4c**). This observation indicates that a higher NAD^+^ pool is a feature of the genetic background, and it is exacerbated during muconate production. The ATP/ADP ratio was 0.82 and 0.73 for KT2440 and 1.30 and 1.52 for CJ781 in the glucose-only and glucose plus 4HB conditions, respectively (**Fig. S11**), similarly showing increased ATP for CJ781 generally, but especially during muconate production. This difference was reflected in 1.3- and 3.1-fold increased total ATP pools in CJ781 as compared to KT2440 on glucose only and glucose plus 4HB, respectively (**Fig. 4d**). These experimental data are consistent with the higher ATP supply predicted by the metabolic flux model.

To further understand the implications of the higher ATP supply, we examined the global changes to ATP-demanding or stress response proteins, as an apparent excess in ATP supply can indicate a cellular response to combat stress.^53,55^ When fed glucose plus 4HB, CJ781 had 103 more proteins significantly elevated that were annotated as involved in transport, degradation, energy, stimulus response, and stress responses compared to KT2440 (**Fig. S9**). Relative to KT2440, CJ781 had 43 ATP-requiring proteins with >2-fold increased abundance (**Fig. 4e**). Incorporation of such ATP-demanding stress responses into the model would likely decrease the apparent excess. Nonetheless, experimental measurements of the ATP pool supported that CJ781 maintained ATP beyond the demands required for anabolism and stress responses.

### Overexpression of the citrate synthase *gltA* mitigates acetate and pyruvate secretions

Accumulation of aliphatic acids – here, acetate and pyruvate for CJ781 (**Fig. 2d**) – in bioproduction schemes could interfere with both downstream muconate separations and cellular growth. Mitigating acetate secretion is complicated by the uncertainty in how acetate is generated in *P. putida* under aerobic growth conditions. There are several putative acetate production pathways, which broadly separate into: 1) direct conversion from acetyl-CoA, 2) byproduct formation from cysteine and ornithine biosythesis, or 3) byproduct formation from other degradation pathways, such as acetaldehyde (**Fig. 5a**).^56^ In addition to generating acetate, the cell can manage excess intracellular acetate by either converting it to acetyl-CoA for reuse or exporting it for future use (**Fig. 5a**).

**Figure 5.**
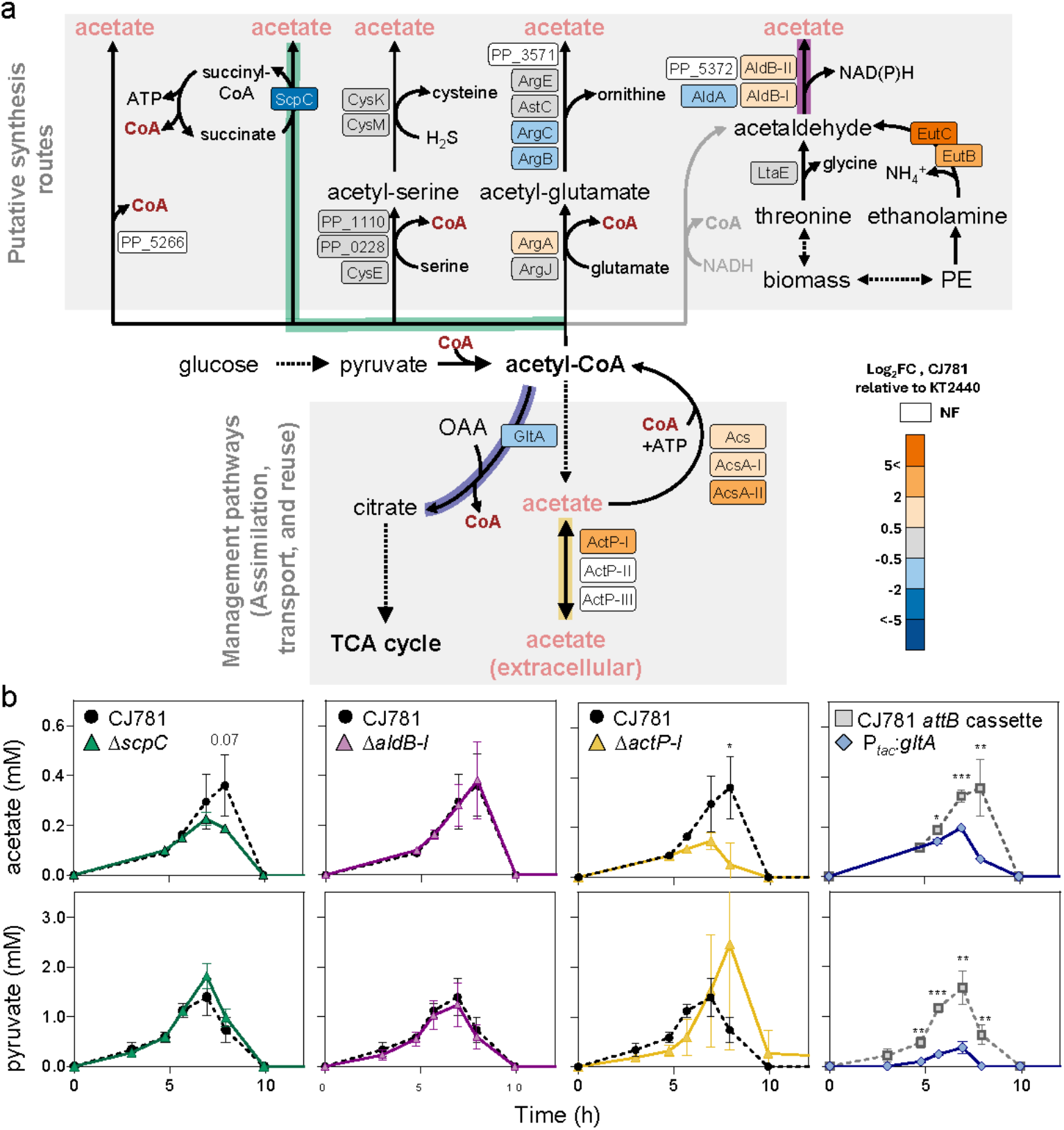
Strategies to control secretion of acetate and pyruvate. (**a**) Schematic of pathways that can produce acetate in *P. putida* paired with differential protein abundances for CJ781 relative to KT2440 when fed 20 mM glucose and 5 mM 4HB. (**b**) Extracellular profiles of engineered strains relative to the control. The black circles represent CJ781, the control for gene knockouts. The grey squares represent RW147 (CJ781 containing *attB* cassette for SAGE integration, which was confirmed to have no altered glucose or 4HB metabolism, **Fig. S14**), the control for the *gltA* overexpression. All data represent the mean ± the standard deviation of three independent biological replicates. Significant differences (p < 0.05) were determined with an unpaired two-tailed t-test between the engineered strains and the control. All raw data are provided in **Excel file 1**.

To probe which acetate pathways could be active in CJ781, we leveraged the differential protein abundance data and genetic engineering in the CJ781 background. The proteomics data indicated that direct conversion from acetyl-CoA was unlikely, as the ATP-independent acetyl-CoA hydrolase (PP_5266) was not found in either dataset, and the acetate:succinate CoA-transferase (ScpC) was 7.0-fold lower for CJ781 relative to KT2440. Fittingly, deleting *scpC* from CJ781 did not significantly decrease acetate secretion (**Fig. 5b**). Proteins in the cysteine and ornithine biosynthesis pathways were unchanged or inconsistent (**Fig. 5a**). Since disruption of amino acid biosynthesis pathways can result in auxotrophy and negative growth effects, we did not delete genes from this pathway. The acetaldehyde dehydrogenases AldB-I and AldB-II were elevated 1.5-fold, but it is also unclear how acetaldehyde would be generated (**Fig. 5a**). Conversion from acetyl-CoA is plausible, but such an enzyme has not been annotated or characterized for KT2440. Formation could alternatively occur via degradation of the amino acid threonine or the phospholipid phosphatidylethanolamine (PE).^57^ PE catabolism is catalyzed by EutB and EutC, which were elevated by 8.4- and 35.2-fold, respectively, in CJ781 relative to KT2440. However, individual deletions of *aldB-I* or *aldA* did not impact acetate secretion (**Fig. 5b, Fig. S12**). Further, the stacked deletion of *aldB-I, aldB-II*, and *eutBC* did not decrease acetate secretion, but increased secretion of glucose oxidation products (**Fig. S13**).

Although these efforts did not determine the production route, we nevertheless sought to reduce secretion by managing the accumulated acetate pool. The three acetyl-CoA synthases (Acs) were elevated by 2.4-to 9.1-fold in CJ781 relative to KT2440 (**Fig. 5a**), indicating acetate is likely reassimilated into central carbon metabolism. Thus, we first considered preventing export of the acetate pool. The acetate permease ActP-I that facilitates acetate import/export was elevated 5.3-fold (**Fig. 5a**). Deletion of *actP-I* in CJ781 decreased the extracellular acetate by 7.2-fold (**Fig. 5b**), demonstrating that deletion of export proteins can mitigate acetate secretion. However, this strategy is not preferred for bioconversion as acetate can often be an additional carbon source, especially in lignin streams.^58,59^

Acetate export instead of reassimilation into acetyl-CoA may be due to a bottleneck for acetyl-CoA entering the TCA cycle. The secretion of pyruvate paired with the 1.9-fold depletion of citrate synthase GltA for CJ781 relative to KT2440 supported this notion (**Fig. 2b**; **Fig. 5a**). The 2.3-fold greater NAD^+^ to NADH pool indicated that this potential bottleneck into the TCA cycle was not due to a buildup in the reduced cofactor and the unavailability of the electron acceptor (**Fig. 3b**). We overexpressed *gltA* in RW147, which is a CJ781-derivative containing an *attB* cassette for serine recombinase–assisted genome engineering (SAGE),^60^ and found that the extracellular concentration of acetate and pyruvate were reduced by 1.7- and 4.2-fold, respectively (**Fig. 5b**). Thus, although it is likely not addressing the root cause of acetate secretion, *gltA* overexpression mitigates acetate and pyruvate accumulation and secretion by promoting acetyl-CoA utilization via the TCA cycle.

### Accumulation of aromatic catabolic intermediates can be overcome by increased expression of *catA2* and *aroY*

CJ781 secreted the aromatic intermediates PCA and catechol at 26.2 ± 9.0% and 16.5 ± 5.9% of the 4HB uptake rate, respectively (**Fig. 2d, Fig. 6a**), indicating substantial bottlenecks in the muconate production pathway. To improve flux from 4HB to muconate, we evaluated the abundances of catabolic pathway proteins. Despite only having deleted *pcaHG* and *catBC* and the regulators *catR* and *pobR* to reroute hydroxybenzoates to muconate in CJ781, all enzymes functioning downstream of PcaHG and CatBC were depleted greater than 16-fold relative to KT2440 (**Fig. 6a**). This difference indicates that the presence of 4HB and PCA in CJ781 did not cause the upregulation of the *ortho*-cleavage pathway genes, and thus the cell avoided investment in unnecessary proteins. Additionally, the native catechol 1,2-dioxygenase, CatA2, was not significantly elevated for CJ781 relative to KT2440 and was even depleted 222-fold relative to the glucose-only conditions (**Fig. 6a**; **Fig. S15**), demonstrating that the introduction of the heterologous pathway bypasses upregulation of *catA2* through *benR*.^61^

**Figure 6.**
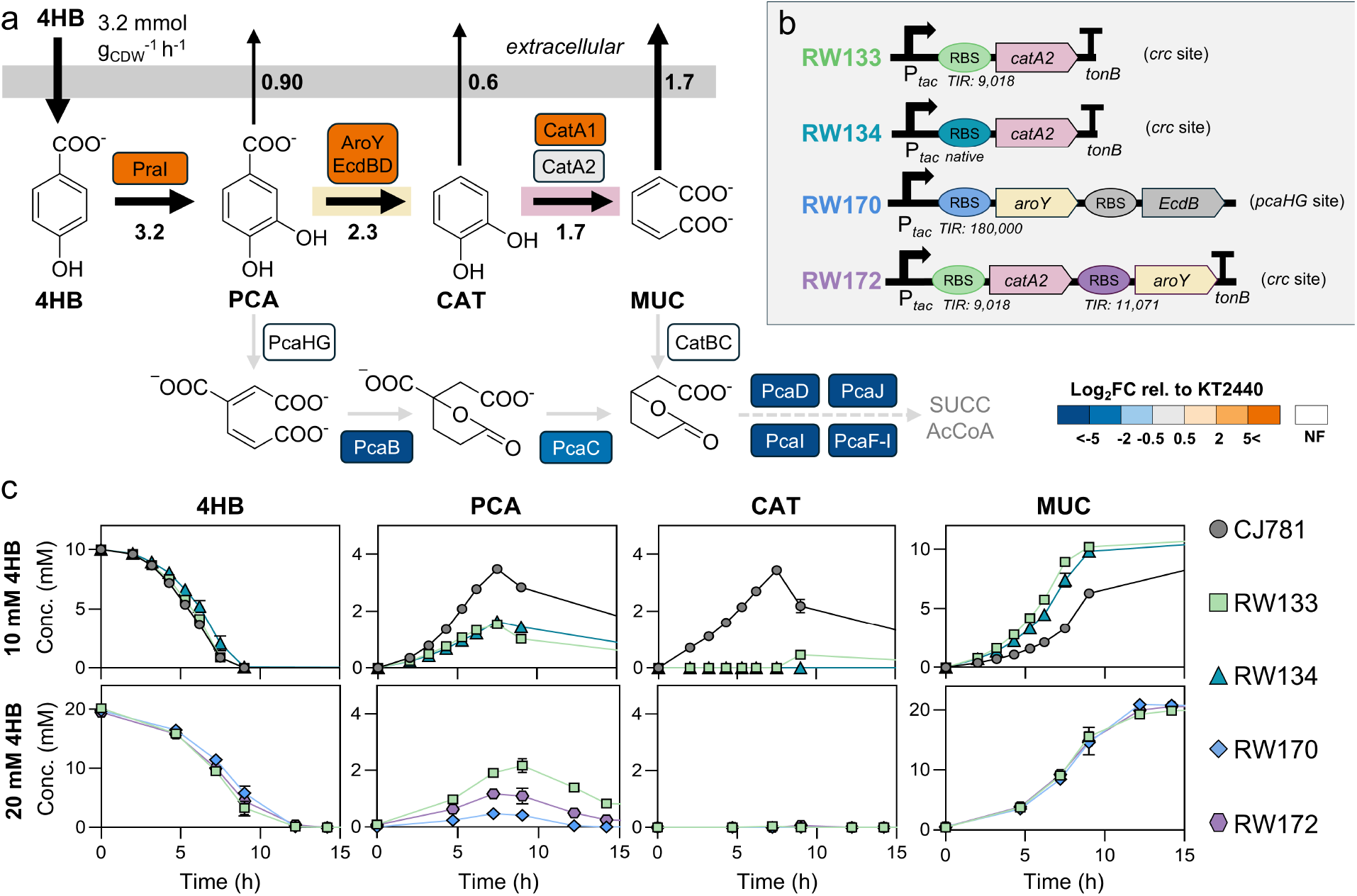
Characterization of engineered strains to improve muconate production. (**a**) Aromatic carbon flux through the engineered muconate production pathway for CJ781 paired with the log_2_ fold-changes (FC) for the measured aromatic catabolic proteins for CJ781 relative KT2440 when fed 20 mM glucose and 5 mM 4HB. (**b**) Schematic of our engineering strategies to improve expression of *catA2* and *aroY* in CJ781. (**c**) Profiles in shake-flask experiments of 4-hydroxybenzoate (4HB), protocatechuate (PCA), catechol (CAT), and muconate (MUC) in the extracellular milieu over time for CJ781, RW133, RW134, RW170, and RW172. Cells were grown on 20 mM glucose with either 10 mM 4HB or 20 mM 4HB. All data in **c** represent the mean ± the standard deviation of three independent biological replicates. All raw data can be found in **Excel file 1**.

To improve flux to muconate, we prioritized first addressing the catechol secretion because a downstream bottleneck can create or exacerbate bottlenecks earlier in the pathway. We overexpressed a second copy of *catA2* at the *crc* deletion site with either a synthetically optimized ribosome binding site (RBS; RW133) or the native RBS (RW134) (**Fig. 6b, Table 1**). Addition of an overexpressed *catA2* into CJ781 did not alter the growth rate (**Fig. S16**), but eliminated catechol secretion, decreased the extracellular PCA by 1.5-fold, and increased the muconate production rate up to 3.0-fold (**Fig. 6c, Fig. S17**). Although RW133 and RW134 performed similarly, the muconate production rate was 20% higher for RW133 (**Fig. S17**).

Although RW133 exhibited lower PCA secretion relative to CJ781, the PCA bottleneck was still observed. To decrease PCA secretion, we either replaced the current RBS in front of *aroY* with one that has a higher translation initiation rate (RW170) or added a second copy of *aroY* into the *catA2* overexpression cassette (RW172) (**Fig. 6b, Table 1**). RW170 and RW172 did not alter the growth rate compared to RW133 (**Fig. S16**), indicating the increased heterologous protein expression did not impact growth. In addition, the extracellular PCA decreased by up to 5.4-fold and 2.0-fold in RW170 and RW172, respectively (**Fig. 6c**). The muconate production rate was improved by 11% for RW170 relative to RW133 (**Fig. S17**). In sum, these data suggest that flux through oxidative catechol ring opening was the primary bottleneck in the muconate production pathway.

### The debottlenecked strains improve titer and rate in bioreactors

We next evaluated the performance of CJ781, RW133, and RW170 in 1-L fed-batch bioreactors to understand the impact of the various genetic modifications on muconate production. 4HB was supplied at a constant feeding rate of 10 mmol h^-1^ normalized to batch volume (L) during the fed-batch phase, together with glucose at a 1:2 molar ratio of glucose to 4HB (see methods). In agreement with flask experiments (**Fig. 6c**), RW133 and RW170 outperformed CJ781 (**Fig. 7a, Fig. S18**). Muconate titer was 3-fold lower for CJ781 than the engineered strains due to early PCA and catechol accumulation which, in turn, triggered 4HB accumulation and ultimately ended the run prematurely at 46.5 h. RW133 and RW170 showed comparable performance across all evaluated metrics and did not accumulate catechol at any point during cultivations (**Fig. 7a, Fig. S18**). RW170 did not accumulate PCA either, and RW133 only accumulated a small amount of PCA (< 0.3 g L^-1^), which did not have a measurable impact on muconate production; however, around 54 h, both strains began to accumulate 4HB, resulting in decreased muconate yields after that time point (**Fig. 7a, Fig. S18**). These results demonstrate that alleviating the catechol bottleneck significantly enhances strain performance. Overall, RW133 achieved a maximum average muconate titer of 46.7 ± 4.3 g L^-1^ at a productivity of 0.9 g L^-1^ h^-1^, while maintaining ∼100% molar yields.

**Figure 7.**
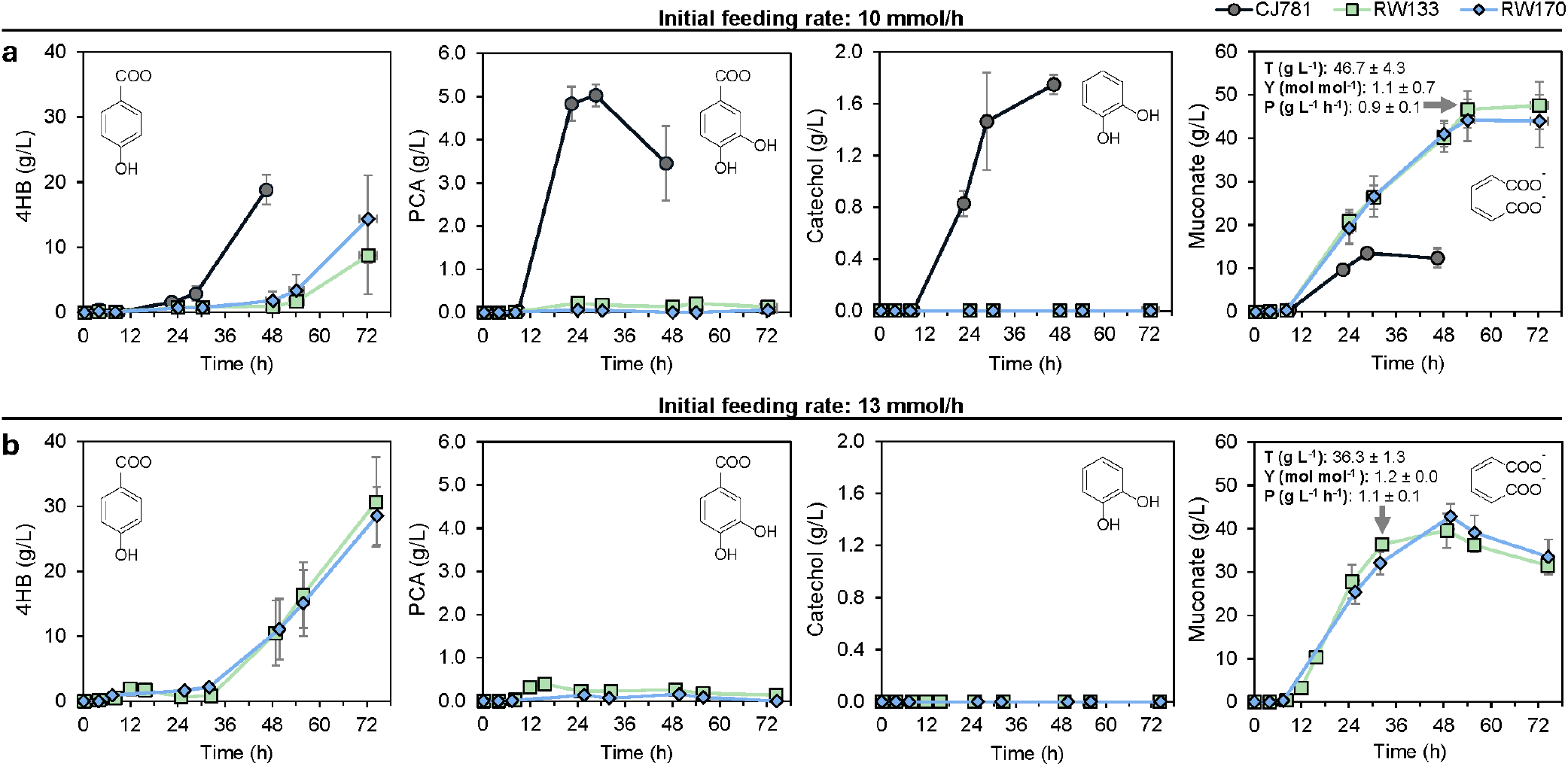
Bioreactor profiles in fed-batch mode for engineered strains RW133 and RW170 compared to base strain (CJ781). 4-hydroxybenzoate (4HB) feeding rates were maintained at (**a**) 10 mmol h^-1^ and (**b**) 13 mmol h^-1^, normalized to the batch volume. 4HB, protocatechuate (PCA), catechol, and muconate concentrations in the supernatant are shown for strains CJ781, RW133, and RW170. Grey arrows indicate the strain and point at which maximum titer (T, g muconate L^-1^), yield (Y, mol muconate mol^-1^ 4HB fed), and productivity (P, g muconate L^-1^ h^-1^) were calculated. These values were calculated before substantial 4HB accumulation, and values are shown alongside the muconate temporal data. Data represent the mean of biological duplicates, except for RW170 at the 10 mmol h^-1^ feeding rate, which was conducted in triplicate. Vertical error bars represent the absolute error between biological duplicates or the standard deviation among triplicates. Horizontal error bars represent the absolute time (h) difference between biological duplicates or the standard deviation among triplicates. Note: The data points corresponding to 12 h and 16 h in RW133 at 13 mmol h^-1^ were obtained from a single replicate.

To investigate the maximum achievable productivities of RW133 and RW170 while aiming to maintain high titers and yields, we increased 4HB feeding rates to 13 and 17 mmol h^-1^. Muconate production remained similar between strains at both feeding rates, although variability among biological triplicates increased at 17 mmol h^-1^ (**Fig. 7b, Fig. S19a**). Catechol accumulation was not observed and PCA accumulation was marginal at these feed rates (reaching a maximum concentration of 0.39 g L^−1^ at 16 h in RW133). However, 4HB began building up at earlier time points than at the 10 mmol h^-1^ feed rate (**Fig. 7b, Fig. S19b-c**). This dynamic with 13 mmol h^-1^ led to lower maximum titers (36.3 ± 1.3 g L^-1^ at 13 mmol h^-1^) but higher productivity (1.1 g L^-1^ h^-1^) prior to 4HB accumulation. These results suggest that 4HB accumulation is not solely due to product or salt buildup but may also reflect an additional metabolic bottleneck in these strains (e.g., transport of aromatic compounds).

Lastly, we evaluated glucose-derived metabolites, including 2-KG, acetate, and pyruvate to better understand carbon metabolism in bioprocess relevant conditions. No consistent trends were observed among strains or feed rates (**Excel File 1**); however, glucose accumulated toward the end of the cultivations (up to 16 g L^−1^), coinciding with 4HB buildup. This accrual was accompanied by 2-KG build up (up to 10 g L^−1^) and acetate production, although the latter generally remained below 0.5 g L^−1^ (**Excel File 1**). Pyruvate was only produced by strain CJ781 up to 0.24 g L^−1^ and in one replicate of RW133 up to 0.20 g L^−1^ (**Excel File 1**). Overall, these results indicate that, in these experimental conditions with controlled feeding of glucose at a 1:2 molar ratio to 4HB, metabolic intermediates do not accumulate until bacterial metabolism slows down at the end of the cultivations.

## DISCUSSION

Producing aromatic catabolic intermediates like muconate from lignin-related aromatic compounds requires supplementation with an alternative carbon source such as glucose or acetate to support growth.^10,15,44–46^ Key to optimizing strains with these partitioned pathways where product formation is decoupled from growth is understanding the metabolic crosstalk between product synthesis and the cellular networks governing energy and biomass generation. In this work, we evaluated the previously engineered muconate producing strain CJ781^10^ by feeding glucose with or without the addition of 4HB, determining rearrangements of the carbon and energy metabolism and identifying metabolic bottlenecks.

Dynamic remodeling of central carbon metabolism in wild-type and engineered *P. putida* KT2440 has been reported previously in response to aromatic compounds,^44,49,62^ catechol stress,^45^ oxidative stress,^63^ and heterologous protein production.^53^ In contrast to catechol as a substrate for muconate production,^45^ heterologous 4HB-to-muconate production in CJ781 triggered increased glucose uptake, elevated periplasmic flux to 2-KG, and increased flux through the oxidative TCA cycle, supporting a higher total production as well as an apparent excess in NADPH and ATP beyond the metabolic demands (**Figs. 2-4**), which more closely resembled flux reorganization from heterologous protein production.^53^ Elevated NADPH production during stress, as demonstrated here, has been characterized as a key trait for *P. putida*,^54,64,65^ and has been observed in *P. putida* and other microorganisms in response to heterologous protein expression.^53,66–68^ Although both heterologous protein expression and catechol stress decreased glucose uptake relative to wild type,^45,53^ cultivations of CJ781 that produced muconate from 4HB significantly increased glucose uptake. Exposure to toluene or styrene similarly increased glucose uptake for *P. putida* DOT-T1E, illustrating a variable response to cellular stressors for *P. putida*.^69^ Overall, these findings underscore the metabolic flexibility of *P. putida* as a platform strain that can maintain cellular fitness and energy balance under different stress conditions.

Although acetate is commonly reported as an overflow mechanism for the regeneration of NAD^+^ in *Escherichia coli*,^70^ acetate leakage has been reported rarely for *P. putida*^71–73^ and the production pathway remains to be elucidated. Previously, *P. putida* KT2440 Δ*crc* grown on complete media secreted pyruvate at about 10 mmol g_CDW_^-1^ h^-1^ and acetate at close to 2 mmol g_CDW_^-1^ h^-1^, which were attributed to metabolite leakage due to the disordered consumption of compounds in LB medium compared to wild type.^72^ We did not observe elevated glucose uptake rate or metabolite accumulation for CJ781 when fed glucose only, indicating that the *crc* deletion alone was insufficient to cause this dysregulation but co-feeding and muconate production triggered this overflow metabolism for CJ781. More recently, acetate formation by electrogenic KT2440 under anaerobic conditions was lessened in the deletion mutants Δ*acsA-I* Δ*acsA-II*, ΔPP_5266, Δ*scpC*, and Δ*aldBI* Δ*aldBII*.^71^ Here, under aerobic conditions, we found that Acs, AcsA-I, and AcsA-II were elevated from 2.4-to 9.1-fold for CJ781 relative to KT2440 (**Fig. 5**), but deletion of these genes would prevent assimilation back to acetyl-CoA and additional biochemical evidence is needed to confirm the reversibility of these reactions. In addition, PP_5266, ScpC, and amino acid biosynthesis did not appear to be involved in acetate production for CJ781 (**Fig. 5**). Likewise, the stacked deletion of *aldB-I, aldB-II*, and *eutBC* did not decrease acetate secretion for CJ781. Additional work is needed to better understand acetate generation in engineered *P. putida* and whether there is a link to membrane degradation during aromatic-to-muconate production.

In lieu of a well-defined route for acetate biosynthesis, we instead focused on strategies to reassimilate intracellular acetate into metabolism. Previously, citrate synthase *gltA* overexpression was used as a strategy to overcome bottlenecks in acetate utilization during the production of succinate.^74^ Here, overexpression of *gltA* mitigated both acetate and pyruvate secretion in shake flask cultivations of CJ781 (**Fig. 5b**). Nonetheless, during bioprocess relevant conditions, pyruvate and acetate accumulated at less than 0.5 g L^-1^ (**Excel file 1**). Thus, bioprocess optimization may offer an alternative to genetic engineering for overcoming metabolite leakage from glucose metabolism. Ultimately, this work underscores the importance of further research to understand the mechanistic differences in metabolism between cultivations in shake flasks and fed-batch cultivations in bioreactors.

Prior work has increased muconate production through improved aromatic catabolism and tolerance.^8,9,28,36,37,39,75^ Although previous *p-*coumarate-fed cultures with CJ781 at 9 mmol h^-1^ achieved maximum titers of 39 g L^-1^ at a productivity of 0.5 g L^-1^ h^-1^,^10^ in this work, 4HB-fed cultures with the same strain at 10 mmol h^-1^ produced muconate at a maximum titer of 13.5 g L^-1^ before accumulating toxic amounts of catechol, PCA, and 4HB (**Fig. 7**). This result demonstrates that feeding 4HB directly, rather than *p*-coumarate, exacerbates bottlenecks in the aromatic-to-muconate pathway. Here, the metabolic bottlenecks at PCA and catechol were alleviated primarily by the addition of a second overexpressed copy of *catA2* and moderately by the tuned expression of *aroY* (**Fig. 6**). This additional engineering did not impact the growth rate of the strains, suggesting that the glucose metabolism mitigated the metabolic burden of increased heterologous protein expression without additional genetic interference. Performance of these new strains when fed 4HB at 13 mmol h^-1^ in bioreactors led to substantial improvement in muconate productivity, reaching up to 1.1 g^-1^ L^-1^ h^-1^ while maintaining a comparably high titer of 36 g L^-1^ to the previous study on CJ781 fed *p*-coumarate.^10^

Employing adaptive laboratory evolution, parallel work by our group demonstrated that mutations in CatA1 mitigated the catechol dioxygenase bottleneck by increasing the abundance of CatA1 and moderately improving its catalytic efficiency to levels comparable to CatA2.^76^ Related work by our group also showed that muconate production from hydrolysate sugars was improved by increasing expression of CatA1 to eliminate catechol accumulation.^77^ Thus, engineering catechol dioxygenase, whether by CatA1 or CatA2 expression, represents a key strategy to debottleneck muconate production. Future work will investigate whether relieving catechol accumulation enables higher feeding rates in bioreactors by enhancing the conversion rates of other aromatic compounds to muconate (e.g., *p*-coumarate, ferulate, and vanillate).

In this work, we used the model lignin-related compound 4HB to mitigate aromatic compound stress and probe the heterologous PCA-to-catechol bypass through AroY to produce muconate.^42^ However, bioprocess relevant feedstocks obtained from plant biomass are more complex and additional work to maintain this high titer, rate, and yield on complex substrate mixtures is needed to ensure the economic viability of this process.^32–35,43,78–80^ Adaptive laboratory evolution experiments and RB-TnSeq represent promising strategies to reveal genetic engineering targets for continued improvements to substrate catabolism and product tolerance.^30,36–39,75^ Overall, this work demonstrated the integration of systems and synthetic biology approaches to uncover metabolic remodeling of glucose metabolism in *P. putida*, which met the cellular energy and redox demands and supported the additional improvements to muconate production from lignin-related aromatic compounds.

## METHODS

### Strain construction

Derivative strains of *P. putida* KT2440 were constructed as outlined previously^10,81–83^ and specific details are provided in **Table S3-S5**. In brief, markerless gene deletions and insertions were performed using pK18sB vector with 1000 base pair homology regions and *sacB/KanR* counterselection.^83^ Overexpression of *gltA* was accomplished by serine recombinase-assisted genome engineering (SAGE) using the pJH0419 vector to target the BxB1 site in RW147 (*P. putida* CJ781 derivative containing BxB1_RV_phi370 cassette).^81,82^ For overexpression targets, ribosome binding sites were optimized using the Salis optimization tool for each gene sequence as described previously.^84^ All plasmids were ordered from TWIST Biosciences or assembled using New England Biolabs HiFi Gibson assembly Mix as described in **Table S4**. Transformants were screened with colony polymerase chain reaction and confirmed with Oxford Nanopore sequencing at Plasmidsaurus. Correct colony transformants were grown overnight at 30 °C in Luria-Bertani (LB) medium (Lennox) and stored at ™80 °C as 20% (v/v) glycerol stocks.

### Media and bacterial cultivations

Cells were cultivated in either LB medium or a modified M9 minimal media (6.78 g L^™1^ Na_2_HPO_4_ 3 g L^™1^ KH_2_PO_4_, 0.5 g L^™1^ NaCl, 1 g L^™1^ NH_4_Cl, 2 mM MgSO_4_, 100 µM CaCl_2_, and 18 µM FeSO_4_) containing 20 mM glucose alone or 20 mM glucose plus varying concentrations of 4HB. Glucose was supplemented into M9 minimal media from a sterile 2 M stock solution to a final concentration of 20 mM. For addition of 4HB, a 100 mM stock solution was prepared by titrating the solution with 5 M NaOH to solubilize and neutralize the aromatic compound in water before adding glucose and M9 minimal media components to prepare a final 4HB concentration of either 5 mM, 10 mM, or 20 mM in 20 mM glucose M9 minimal media. For the ^13^C-tracer experiments, media was prepared with 5 mM unlabeled 4HB and 20 mM glucose which was either 100% [1-^13^C]-glucose, 100% [6-^13^C]-glucose, or an equimolar mixture (50:50) of [U-^13^C_6_]-glucose and unlabeled glucose. Before using media for bacterial cultivation, all media was filter sterilized (0.2 μm pore size). Chemicals for media components were purchased from Millipore Sigma. Isotope tracers were purchased from Cambridge Isotopes.

For all experiments, glycerol stocks were streaked onto LB agar plates and grown overnight at 30 °C. Individual colonies, in triplicate, were selected and inoculated into 14-mL round bottom Falcon^®^ tubes containing 5 mL of LB medium and incubated for 12-15 h at 30 °C and 225 rpm. These LB seed cultures were washed twice with M9 salts (6.78 g L^™1^ Na_2_HPO_4_, 3 g L^™1^ KH_2_PO_4_, 0.5 g L^™1^ NaCl, 1 g L^™1^ NH_4_Cl) and then inoculated into 25 mL or 5 mL of experimental media (M9 minimal media containing glucose only or glucose plus 4HB) either in 125-mL baffled flasks or in 14 mL tubes to an initial OD_600_ of 0.07. These second seed cultures were transferred again into 25 mL of the same experimental media in 125-mL baffled flasks to an initial OD_600_ of 0.07 for experimental measurements. For cultures containing 20 mM glucose with 20 mM 4HB, the pH values of the flasks were monitored at each sampling time point using a mini pH meter (HORIBA LAQUAtwin pH-33) and adjusted to a pH of 7 with 1 N NaOH. Additionally, at 12 h, these cultures were spiked with sterile 2 M glucose to a final concentration of 20 mM to ensure enough glucose was present to support the conversion of 20 mM 4HB to muconate. All cultures were grown at 30 °C with shaking (225 rpm).

### Physiological characterizations

Aliquots of cell suspensions were collected over time to monitor growth at an optical density of 600 nm (OD_600_) and to quantify extracellular metabolites. Samples for metabolite quantification were collected by centrifuging for 5 min at >10,000*g* followed by filtering the supernatant through 0.22 µm nylon Costar® Spin-X® centrifuge tube filters (Corning). All filtered extracellular samples were stored at -20 °C until analysis. For determining the OD to g_CDW_ conversion factors, the cell pellet aliquots across exponential growth were stored at -20 °C overnight before lyophilizing for 24 h and weighing the dry biomass. Growth rates were determined by fitting the exponential timepoints to the growth model described previously.^85^ The OD to g_CDW_ conversion factor was determined for each strain by regression analysis of the linear fit between OD and g_CDW_ (R^2^ coefficient greater than 0.96) (**Table S1**). A linear fit was also used to determine consumption and secretion rates based on the quantified extracellular profiles of substrate depletion and metabolite production over time. These rates were then normalized to growth rate and g_CDW_.

### Assaying NAD^+^/NADH and ATP/ADP pools

Cells of KT2440 and CJ781 were cultured and transferred into 20 mM glucose alone or 20 mM glucose plus 5 mM 4HB as described above. During exponential growth (OD_600_ ∼ 0.5), three technical replicates were harvested from each of three independent shake flasks for a total of nine 1 mL samples per strain and growth condition. Each sample was aliquoted for OD_600_ measurements, the ATP/ADP assay (MAK135, Millipore Sigma), and the NAD^**+**^/NADH assay (MAK468, Millipore Sigma). For both assays, cell suspensions were pelleted via centrifugation at 5,000 *g* for 10 min at room temperature and then washed with phosphate buffered saline (PBS). For the ATP/ADP assay, the washed cell pellet was resuspended in an equal volume of PBS, whereas for the NAD^**+**^/NADH assay the washed cell pellets were resuspended in the supplied extraction buffers. For both assays, extractions and measurements were performed according to the manufacturer’s instructions.

### Intracellular metabolite extractions

During exponential growth (OD_600_ of ∼0.4-0.5), cells were extracted via rapid filtration and quenching as described previously.^86^ Timepoints were chosen to occur during the co-depletion of glucose and 4HB for each strain (**Fig. S1)**. In brief, cells were filtered onto 0.45 µm filter discs and extracted with a solution of cold (-20 °C) 2:2:1 methanol:acetonitrile:water. The lysed cell debris were pelleted via centrifugation at >15,000 *g* for 1 min at 4 °C. The supernatant of the extract was stored immediately at -80 °C or dried under N_2_ gas before storage at -80 °C until HPLC-MS analysis.

### Intracellular and extracellular metabolomics analysis via UHPLC-MS

Metabolomics analysis of intracellular metabolite extracts and extracellular milieu was conducted via liquid chromatography mass spectrometry (LC-MS) as described previously.^86^ In brief, a Vanquish ultra-high-performance liquid chromatography (UHPLC) system (Thermo Scientific) was coupled to a hybrid quadrupole-Orbitrap™ mass spectrometer (Q Exactive™; Thermo Scientific) equipped with electrospray ionization operating in negative-ion mode. A 2.1 × 100 mm reverse-phase C_18_ column with a 1.7 μm particle size (Water™; Acquity UHPLC BEH) was used for chromatography and maintained at 25 °C. Solvent A consisted of 97:3 H_2_O: methanol + 10 mM tributylamine and Solvent B consisted of 100% methanol. Two chromatography gradients were used for analysis. The first was as follows: 0–2.5 min, 5% B; 2.5–17 min, linear gradient from 5% B to 95% B; 17–19.5 min, 95% B; 19.5– 20 min, linear gradient from 95% B to 5% B; 20–25 min, 5% B. The second gradient was as follows: 0–2.5 min, 5% B; 2.5–7.5 min, linear gradient from 5% B to 20% B; 7.5–13 min, 20% B to 55% B; 13–18.5min, 55% B to 95% B; 18.5–19 min linear gradient from 95% B to 5% B; 19–25 min, 5% B. For both gradients, the flow rate was held constant at 0.2 mL min^-1^. Analytes of interest were identified by retention times based on pure standards and monoisotopic mass using Metabolomic Analysis and Visualization Engine (MAVEN)^87^ and El-MAVEN^88^ software. The natural abundance of ^13^C in all isotopologue data was corrected manually. Quantification of analytes was obtained using commercial standards (Millipore-Sigma or Fisher Scientific) with an R^2^ coefficient of 0.992 or higher for the calibration curve.

### Analysis of aliphatic organic acids by HPLC-DAD-RID

Quantitation of aliphatic organic acids—gluconic, 2-ketogluconic, pyruvic, acetic, lactic, and succinic acids—was carried out using high-performance liquid chromatography equipped with refractive index and diode-array detection. Separation was achieved on a Rezex ROA-Organic Acid column with 0.02 N sulfuric acid (0.01 M) as the mobile phase, following the method described previously (xx) ^89^with minor modifications, detailed below. Gluconic, 2-ketogluconic, and pyruvic acids were included as additional analytes; their stock and calibration standards were prepared identically to those of the other acids in the protocol. To ensure accurate quantitation and minimize co-elution with matrix components, diode-array detection at 210 nm was used for gluconic, 2-ketogluconic, and pyruvic acids, while refractive index detection remained in use for the remaining acids.

### Quantification of 4-hydroxybenzoic acid, muconic acid isomers, catechol, and protocatechuic using UHPLC-DAD

Analysis of the *cis,cis* (c,c) and *cis,trans* (c,t) isomers of muconic acid, along with aromatic compounds, was performed using an ultra-high-pressure liquid chromatography (UHPLC) system as described previously.^90^ Briefly, samples were analyzed by UHPLC equipped with diode-array detection. Chromatographic separation was achieved using an Acquity BEH C18 column with a mobile-phase gradient consisting of 0.2% formic acid in water and acetonitrile.

### Glucose analysis via HPLC-ELSD

Due to co-elution of gluconic acid in other analytical methods, glucose was quantified using high-performance liquid chromatography with evaporative light-scattering detection (HPLC-ELSD) as described previously.^91^ Briefly, chromatographic separation of glucose and other supernatant constituents was achieved using a Shodex SUGAR SZ5532 column and a mobile-phase gradient consisting of (A) 0.1% formic acid in water and (B) 0.1% formic acid in acetonitrile. Detection and quantification of glucose were performed using ELSD, with a calibration based on a quadratic regression curve exhibiting an *r*^2^value ≥ 0.995.

### Proteomics analysis

*Pseudomonas putida* cells were prepared for proteomic analysis following the procedure outlined previously.^92,93^ Briefly, cells were disrupted using bead beating with 0.15 mm zirconium oxide beads in 100 mM Tris-HCl buffer at pH 8.0, using a Geno/Grinder 2010 (SPEX). The resulting lysates were brought to a final concentration of 4% SDS and 10 mM dithiothreitol, then heated at 90 °C and clarified by centrifugation. Protein concentration was determined using a NanoDrop One^C^ spectrophotometer at 205 nm after diluting the samples 20-fold in water. The supernatant was transferred to fresh tubes, and free cysteines were alkylated with 30 mM iodoacetamide. Proteins were then isolated using the protein aggregation capture (PAC) method,^94^ washed according to protocol, and digested *in situ* with sequencing-grade trypsin at a 1:75 enzyme-to-protein ratio (w/w) in 100 mM ammonium bicarbonate (pH 8.0). Digestion was carried out at 37 °C overnight and repeated the next day for 3 h with another addition of trypsin. The resulting peptides were acidified with formic acid to 0.5%, filtered using a 10 kDa molecular weight cut-off spin filter (Vivaspin500; Sartorius), and quantified again using a NanoDrop spectrophotometer. For each sample, 3 µg of peptides were subjected to one-dimensional LC-MS/MS analysis using a Vanquish UHPLC system coupled to a nanoelectrospray source interfaced with a Q Exactive Plus mass spectrometer (Thermo Scientific), as previously reported.^93^

Peptide fragmentation (MS/MS) data were searched against the KT2440 and CJ781 proteome, supplemented with common contaminant sequences, using the SEQUEST HT algorithm in Proteome Discoverer v2.5 (Thermo Scientific). Peptide-spectrum matches (PSMs) were evaluated and filtered using Percolator to maintain a false discovery rate (FDR) below 1% at both the PSM and peptide levels. Quantification was based on the chromatographic area under the curve, with peptide signals aggregated to derive protein-level abundance estimates. Statistical processing was performed as outlined previously^92^

### ^13^C-metabolic flux analysis

We reconstructed a core metabolic network for *P. putida* using the genome-scale metabolic model iJN1463^56^ as a basis and added reactions responsible for 4HB metabolism. The biomass reaction was adjusted to account for metabolites in lumped pathways by representing them as their relevant precursors. We then constructed an atom mapping model (AMM) for this network, following information from the AMM of other organisms including *Escherichia coli*^95^ and *Synechococcus* sp.^96,97^ Mappings for the remaining reactions were determined manually. All carbon atoms are represented following a carbon numbering scheme consistent with the IUPAC convention. Both model and AMM are provided in **Excel file 1**.

We used a ^13^C metabolic flux analysis (^13^C-MFA) procedure.^95^ Briefly, this method performs nonlinear optimization via the fmincon function from the optimization toolbox of MATLAB to find a flux solution that minimizes the weighted sum of squared residuals (SSR) between the measured and simulated ^13^C mass distribution vectors (MDVs) and observed uptake and secretion fluxes. The simulated MDVs are determined from the substrate labeling and predicted fluxes and the method employs the elementary metabolite unit (EMU) framework.^98^ ^13^C-MFA also used Gurobi version 11 (https://www.gurobi.com) during confidence interval computations. Python 3.10 was used to read the labeling measurements, compile the measurements for the selected strain and conditions, including measured growth rate and metabolite uptake and secretion rates collected as part of this study, scale the glucose uptake to 100 mmol of glucose uptake g_CDW_^-1^ h^-1^ as the basis, propagate glucose uptake error, and output processed data files for each corresponding strain and condition pair. The complete model and AMM were processed to remove reactions irrelevant to the strain and condition pair, such as reactions eliminated through gene knockouts or 4HB pathways when it was not present in the growth media. This processing pipeline generated consolidated model and AMM files that included dilution reactions for all selected measurements.

The best-fit flux solution was selected from 500 random initialization simulations by selecting the one with the lowest weighted SSR. We performed a χ^2^ goodness-of-fit test for each best-fit solution^99^, and 95% confidence intervals for each flux were estimated as described previously.^99,100^ Solution recovery was determined by counting all solutions from the 500 random initialization that have SSR values within the χ^2^ statistic for a p-value of 0.05 and one degree of freedom (i.e., 3.841) of the SSR of the best-fit-solution. All flux values were then rescaled to the glucose uptake. Tabulated residuals were examined by plotting using matplotlib in Python.

For calculating the cofactor and ATP balances, the fluxes corresponding to reactions that supplied or used cofactors or ATP were summed (**Excel file** 1). The NADPH supply was greater than the demand, and thus all reactions with cofactor flexibility for NADPH or NADH were modeled as supplying NADH. All additional surplus of NADPH was assumed to be converted to NADH through transhydrogenase reactions to supplement energy production. The ATP production from NADH and FADH_2_/UQH_2_ was calculated using a phosphate-to-oxygen ratio of 1.875 and 1.0, respectively

### Bioreactor cultivations in fed-batch mode

To test strains CJ781, RW133, and RW170 in bioreactors, independent seed cultures for each bioreactor were prepared from glycerol stocks in 250 mL baffled flasks containing 50 mL of LB Miller. These seed cultures were incubated at 30 °C and 225 rpm for ∼16-18 h before pelleting the cell biomass through centrifugation at 5,000 *g* for 8 min and resuspending it in 5 mL of M9 medium (13.56 g/L Na_2_HPO_4_, 6 g/L KH_2_PO_4_, 1 g/L NaCl, and 2.25 g/L (NH_4_)_2_SO_4_). Each seed was used to inoculate bioreactors to a target of optical density at 600 nm (OD_600_) of 0.2 (measured with the NanoDrop One, Thermo Scientific).

The bioreactors (1-L my-Control, Applikon Bio) contained an initial batch volume of 250 mL. The medium in the batch phase contained 13.56 g/L Na_2_HPO_4_, 6 g/L KH_2_PO_4_, 1 g/L NaCl, and 2.25 g/L (NH_4_)_2_SO_4_, 0.24 g/L MgSO_4_ x 7 H_2_O, 11 mg/L CaCl_2_ x 2 H_2_O, and 2.73 mg/L FeSO_4_ x 7 H_2_O as well as 2.7 g glucose, the latter utilized as the sole carbon source. The bioreactors were maintained at 30 °C and pH 7 (controlled by 4N NaOH) and air was sparged at 1 vvm. The initial agitation speed and dissolved oxygen (DO) of the batch phase were 350 rpm and 100%, respectively. Once DO dropped to 30%, an automatic agitation cascade was employed to maintain it at this level.

After 4 h of batch growth, a spike of 4HB stock (750 mM) solution was added to a final concentration of 2 mM in the media to induce the expression of enzymes related to aromatic catabolism. The fed-batch phase initiated when glucose was depleted in the batch phase (∼8-10 hours after inoculation), which was indicated by a rapid increase of DO levels. The feed medium consisted of 250 mL of 750 mM 4HB, 370 mM glucose, and 100 mM (NH_4_)_2_SO_4_ titrated to a final pH of 7.7 with 10 N NaOH. Antifoam (4 mL/L) was also added independently to each feeding source. External pumps (Watson Marlow 120U/DV analogue control variable speed pump) were used to deliver the feed, using I/S 13 MasterFlex tubing with a section of I/14 Pharmed tubing positioned in the pump head to improve feed accuracy. The initial feeding rate for the evaluation of strains RW133, RW170, and CJ781 was 3.33 mL/h, which corresponded to 10 mmol/h of 4HBA (normalized by the batch volume). Higher feeding rates were also tested in RW133 and RW170 cultivations, which corresponded to 4.33 and 5.67 mL/h for 13 and 17 mmol/h feeding rates, respectively. Cultivations were terminated when agitation reached minimum values (350 rpm) and glucose began to accumulate. Samples were taken periodically from the bioreactors to evaluate bacterial growth (OD600) and to analyze extracellular metabolites (glucose, 2-ketogluconate, acetate, muconate, protocatechuate, and 4HB).

## Data availability

Experimental data are provided in the manuscript, supplemental materials, or the **Excel file 1**. MFA code and files are available at github.com/maranasgroup/P_putida_MFA. Proteome raw data and search results were deposited in the MassIVE and ProteomeXchange repositories (link: ftp://massive-ftp.ucsd.edu/v11/MSV000099761/) under the accessions MSV000099761 and PXD070362 respectively).

## ACKNOWLEDGEMENTS

This work was authored in part by the National Laboratory of the Rockies for the U.S. Department of Energy (DOE), operated under Contract No. DE-AC36-08GO28308. This material in part is based upon work supported in part by the Center for Bioenergy Innovation (CBI), U.S. DOE, Office of Science, Biological and Environmental Research Program under Award Number ERKP886. Funding was provided in part by the U.S. DOE, Office of Critical Minerals and Energy Innovation, Alternative Fuels and Feedstocks Office. Computations for this research were performed on the Pennsylvania State University’s Roar Collab supercomputer. The authors of this work recognize the Penn State Institute for Computational and Data Sciences (RRID:SCR_025154) for providing access to computational research infrastructure within the Roar Core Facility (RRID: SCR_026424). We thank Hannah Alt for analytical support. The views expressed in the article do not necessarily represent the views of the DOE or the U.S. Government. The U.S. Government retains and the publisher, by accepting the article for publication, acknowledges that the U.S. Government retains a nonexclusive, paid-up, irrevocable, worldwide license to publish or reproduce the published form of this work, or allow others to do so, for U.S. Government purposes.

## Contributions

R.A.W., A.J.B., C.D.M., D.A.N., G.T.B., and A.Z.W. developed the project concept. R.A.W., A.J.B., D.S., G.T.B., and A.Z.W. designed experiments. R.A.W. and M.A.G. performed strain engineering. R.A.W., A.J.B, M.A.G. and A.Z.W. performed shake-flask experiments. R.A.W, A.J.B., M.M.C, E.T., and A.Z.W. generated metabolomics data. R.A.W, R.J.G., D.L.C., and A.Z.W. generated the proteomics data. P.F.S. and J.I.H. performed the ^13^C-metabolic flux analysis. A.F.B. and K.J.R. quantified analytes in extracellular milieu. A.N.M. performed bioreactor cultivations. D.S., R.L.H., C.M., D.A.N., G.T.B., and A.Z.W. provided supervision and funding. R.A.W., D.S., G.T.B., and A.Z.W. wrote the manuscript, with all authors editing and approving the final draft.

## Ethics declaration

Competing interests None to declare.

## Notes

### Competing Interest Statement

The authors have declared no competing interest.

