## Supplemental Information for "Engineered *Pseudomonas putida* reconfigures metabolic fluxes to support energy demands during muconate bioproduction from lignin-related aromatics"

**Table S1.** Characteristics of *P. putida* KT2440 and CJ781. Specific rates for growth, uptake, and secretion calculated during exponential growth phase. In addition, the yields for biomass ($Y_{X/Glucose}$) and the conversion between OD600 and $g_{CDW}$ are listed. Data represent mean ± standard deviation from six independent biological replicates for growth rate and three independent biological replicates for all other rates. NA, not applicable; NF, not found.

| Parameter | Glucose only | | Glucose + 4HB | |
|---|---|---|---|---|
| | KT2440 | CJ781 | KT2440 | CJ781 |
| $\mu_{max}$ ($h^{-1}$) | 0.54 ± 0.03 | 0.54 ± 0.08 | 0.49 ± 0.06 | 0.50 ± 0.08 |
| $Y_{X/Glucose}$ ($g\ g^{-1}$) | 0.30 ± 0.01 | 0.37 ± 0.03 | 0.38 ± 0.07 | 0.23 ± 0.01 |
| Conversion ($g_{CDW}\ L^{-1}\ OD_{600}^{-1}$) | 0.60 ± 0.02 | 0.85 ± 0.13 | 0.71 ± 0.02 | 0.66 ± 0.03 |
| Glucose uptake (mmol $g_{CDW}^{-1}\ h^{-1}$) | 10.07 ± 0.45 | 8.12 ± 0.59 | 7.41 ± 1.33 | 12.15 ± 0.50 |
| 4HB uptake (mmol $g_{CDW}^{-1}\ h^{-1}$) | NA | NA | 3.54 ± 0.70 | 3.20 ± 0.20 |
| GLCN formation (mmol $g_{CDW}^{-1}\ h^{-1}$) | 0.76 ± 0.29 | 0.29 ± 0.19 | 0.31 ± 0.29 | NF |
| 2-KG formation (mmol $g_{CDW}^{-1}\ h^{-1}$) | 1.18 ± 1.30 | NF | 1.46 ± 0.51 | 1.78 ± 0.37 |
| PYR formation (mmol $g_{CDW}^{-1}\ h^{-1}$) | 0.021 ± 0.003 | 0.043 ± 0.020 | 0.057 ± 0.002 | 0.327 ± 0.087 |
| ACE formation (mmol $g_{CDW}^{-1}\ h^{-1}$) | NF | NF | NF | 0.274 ± 0.036 |
| LAC formation (mmol $g_{CDW}^{-1}\ h^{-1}$) | 0.037 ± 0.008 | 0.054 ± 0.007 | 0.049 ± 0.005 | 0.083 ± 0.010 |
| SUCC formation (mmol $g_{CDW}^{-1}\ h^{-1}$) | 0.036 ± 0.006 | 0.038 ± 0.009 | 0.025 ± 0.019 | 0.074 ± 0.008 |
| PCA formation (mmol $g_{CDW}^{-1}\ h^{-1}$) | NA | NA | NF | 0.84 ± 0.29 |
| CA formation (mmol $g_{CDW}^{-1}\ h^{-1}$) | NA | NA | NF | 0.53 ± 0.22 |
| MA formation (mmol $g_{CDW}^{-1}\ h^{-1}$) | NA | NA | NF | 1.62 ± 0.16 |

**Table S2.** Significance parameters of $^{13}C$-metabolic flux analysis.

| Condition | Glucose only | | Glucose + 4HB | |
|---|---|---|---|---|
| Strain | KT2440 | CJ781 | KT2440 | CJ781 |
| DOF | 185 | 183 | 189 | 192 |
| $\chi^2_{1-0.05/2}$(DOF) | 224.56 | 222.35 | 228.96 | 232.27 |
| min opt [SSR] | 143.95 | 202.89 | 136.35 | 184.58 |
| recovery count | 307 | 410 | 404 | 364 |

**Table S3.** Oligonucleotides utilized in this study.

| Name | Sequence (5'->3') | Purpose |
|---|---|---|
| oRW131 | CCATGTGATAGGTACTGGTCGC | Colony PCR of pRW040, pRW041, and pRW060 transformations |
| oRW132 | TTACTGACGGGTGGCTACTG | |
| oRW157 | AAGGCCATTGGGGCTGCATTG | Extract plasmid backbone for pRW060 |
| oRW158 | GAATTCGTAATCATGTCATAGCTGTTTCCTGTGTG | |
| oRW159 | TATGACATGATTACGAATTCGAGCTGTTGACAATTAATCATCGGC | Extract aroY HR for pRW060 |
| oRW160 | TGTTATGTTAATCGGGATATTCAGGCCTCCTGCAAAG | |
| oRW161 | CTTTGCAGGAGGCCTGAATATCCCGATTAACATAACATCCCACTTAGGAAGACTTTTTATGCAGAACCCGATCAACGA | Extract aroY for pRW060 |
| oRW162 | CAATGCAGCCCCAATGGCCTTAGTAAAAAGCCTCCGGTCGGAGGCTTTTGACTTCACTTCTTGTCGCTGAACAG | |
| oRW163 | AGTTTAGACTTCTTGGTCGCGG | Colony PCR of pRW061 integration for screening |
| oRW164 | GCAGATGGTCTCACTTGGTC | |
| oRW165 | GTACACGTTCTGCAGGAAGC | Colony PCR of pRW061 transformation; paired with oRW164 |
| oRW201 | GCGGAAGGAAGAGCTCAC | Colony PCR of pRW076 transformation |
| oRW202 | CTACAGTGCCTTCACCCTG | |
| oRW203 | AAGGCCAGCAGTACCAGC | Colony PCR of pRW077 transformation |
| oRW204 | TACCCAGCAGGTTCAGTGC | |
| oRW205 | GATTTCCTCGTTCAGCGGTG | Colony PCR of pRW078 transformation |
| oRW206 | GTGTCGCTCTGACTGGCTC | |
| oRW207 | CCAAGATCGGCCTGATCGTC | Colony PCR of pRW079 transformation |
| oRW208 | TCGAGGCGACATTGATAATGC | |
| oTLH040 | GTCTTTATAGCTTCGACGTCATAGTAGG | Colony PCR of pRW081 transformation |
| oTLH041 | GAACAGCAGGTTGGTCTTGATGC | |
| oMG061 | GATCTACAACGACGCCAGCCG | |

| | | |
|---|---|---|
| oMG062 | GCAGCCGGCACCAAAGATG | Colony PCR of pIAR007 transformation |
| oMG084 | GCTGTCGCCCAGGATTGCAG | Colony PCR of pMG015 transformation |
| oMG085 | CGCTGTTCGGTATCGGCGGC | |

**Table S4.** Plasmids utilized in this study. TIR, translation initiation rate. RBS, ribosome binding site.

| Name | Description | Construction details | Used to construct | Reference |
|---|---|---|---|---|
| pK18sB | Backbone for gene deletion or replacement via electroporation | Genbank: MH166772. | Backbone for pRW040, pRW041, pRW060, pRW061, and pRW076-79 | Previously described in Jayakody, et al., 2018.2 |
| pGW97 | Used to insert the attB sites for BxB1, RV, and Φ370 at PP_4740 (hsdR). | Plasmid obtained courtesy of Adam Guss' group at Oak Ridge National Laboratory. | RW147 | ([12]) |
| pJH0419 | Sage integrase plasmid for integration in the BxbI site conferring kanamycin resistance | Plasmid obtained courtesy of Adam Guss' group at Oak Ridge National Laboratory. | Backbone for pRW081 | ([12]) |
| pGW031 | Integrase Expression Suicide Vector for integration at the BxBI site | Plasmid obtained courtesy of Adam Guss' group at Oak Ridge National Laboratory. Co-transformed alongside cargo plasmid (pJH0419) | RW249 | ([12]) |
| pRW040 | pK18sB-based plasmid for insertion of $P_{tac}$:rbs$_{synthetic}$:*catA2*:*tonB* in *P. putida* KT2440-derived strains | 1 kb homology regions upstream and downstream of *crc* were designed to insert the expression cassette at the *crc* knockout location. The expression cassette contained $P_{tac}$, RBS (TIR = 9,018), *catA2*, and *tonB*. The RBS was optimized using the Salis optimization tool as described in the methods. An XbaI site was inserted between the two homology regions, and the insert was cloned into the pK18sb backbone at the EcoRI and HindIII sites. The plasmid was synthesized and sequence-verified by Twist Biosciences. | RW133 | This study |
| pRW041 | pK18sB-based plasmid for insertion of $P_{tac}$:rbs$_{NATIVE}$:*catA2*:*tonB* in | 1 kb homology regions upstream and downstream of *crc* were designed to insert the expression cassette at the *crc* | RW134 | This study |

| | | | | |
|---|---|---|---|---|
| | *P. putida* KT2440-derived strains | knockout location. The expression cassette contained $P_{tac}$, native RBS, *catA2*, and *tonB*. The RBS was optimized using the Salis optimization tool as described in the methods. An XbaI site was inserted between the two homology regions, and the insert was cloned into the pK18sb backbone at the EcoRI and HindIII sites. The plasmid was synthesized and sequence-verified by Twist Biosciences. | | |
| pRW060 | pK18sB-based plasmid for insertion of *aroY* into Δcrc::Ptac:catA2 expression cassette in RW133 | 1 kb homology regions were designed to insert RBS and *aroY* into the expression cassette for *catA2.* The RBS (TIR: 11071.26) was optimized using the Salis optimization tool as described in the methods. This plasmid was constructed using Gibson assembly of fragments generated using oRW159, oRW160, oRW161, and oRW162. The plasmid was sequenced by Oxford Nanopore Sequencing. | RW172 | This study |
| pRW061 | pK18sB-based plasmid for replacement of the RBS in front of *aroY* in CJ781 | 1 kb homology regions were designed upstream and downstream of the RBS for *aroY* in CJ781. The RBS (TIR: 179993.62) was optimized using the Salis optimization tool as described in the methods. An XbaI site was inserted between the two homology regions, and the insert was cloned into the pK18sb backbone at the EcoRI and HindIII sites. The plasmid was synthesized and sequence-verified by Twist Biosciences. | RW170 | This study |
| pRW076 | pK18sB-based plasmid for deletion of *scpC* in *P. putida* KT2440-derived strains | 1 kb homology regions upstream and downstream of *scpC* were designed. An XbaI site was inserted between the two homology regions, and the insert was cloned into the pK18sb backbone at the EcoRI and HindIII sites. The plasmid was synthesized and sequence-verified by Twist Biosciences. | RW243 | This study |
| pRW077 | pK18sB-based plasmid for deletion of *aldB-I* in *P. putida* KT2440-derived strains | 1 kb homology regions upstream and downstream of *aldB-I* were designed. An XbaI site was inserted between the two homology regions, and the | RW236 | This study |

| | | | | |
|---|---|---|---|---|
| | | insert was cloned into the pK18sb backbone at the EcoRI and HindIII sites. The plasmid was synthesized and sequence-verified by Twist Biosciences. | | |
| pRW078 | pK18sB-based plasmid for deletion of *aldA* in *P. putida* KT2440-derived strains | 1 kb homology regions upstream and downstream of *aldA* were designed. An XbaI site was inserted between the two homology regions, and the insert was cloned into the pK18sb backbone at the EcoRI and HindIII sites. The plasmid was synthesized and sequence-verified by Twist Biosciences. | RW244 | This study |
| pRW079 | pK18sB-based plasmid for deletion of *actP-I* in *P. putida* KT2440-derived strains | 1 kb homology regions upstream and downstream of *actP-I* were designed. An XbaI site was inserted between the two homology regions, and the insert was cloned into the pK18sb backbone at the EcoRI and HindIII sites. The plasmid was synthesized and sequence-verified by Twist Biosciences. | RW245 | This study |
| pRW081 | pJH0419-based plasmid for the expression of $P_{tac}$:*gltA* | The expression cassette included an operon containing $P_{tac}$, RBS, and *gltA*. The RBS (TIR=9997) was optimized using the Salis optimization tool as described in the methods. This plasmid will integrate into the Bxb1 site. The plasmid was synthesized and sequence-verified by Twist Biosciences. | RW249 | This study |
| pIAR007 | pK18sB-based vector for deletion of aldB-II (PP_2680) in *P. putida* | A 746 bp homology region upstream and a 745 bp homology region downstream of *aldB-II* were designed. The upstream targeting region was amplified with primer pair oIAR015/oIAR016 and the downstream region was amplified with oIAR017/oIAR018 from *P. putida* KT2440 gDNA. pk18sb was digested with EcoRI and HindIII. The pieces were asssembled using Hifi DNA assembly and the assembled plasmid was transformed into NEB 5-alpha F'IQ *E. coli* cells. | MG054 | n/a |
| pMG015 | pK18msBI-based vector for deletion of eutBC (PP_0542 and PP_0543) in *P. putida* | 1kb homology regions upstream and downstream of *eutBC* were designed. This insert was cloned into the pK18msBI | MG075 | This study |

| | | |
|---|---|---|
| | | backbone. The plasmid was synthesized and sequence-verified by Twist Biosciences. |

**Table S5.** Sequences of synthetically optimized ribosome binding sites (RBS) used in this study.

| **Strain** | **Sequence** |
|---|---|
| RW133 | GCTCCCCAATAGCGATCGAGACTTTTAT |
| RW170 | AAAAAACCTCCTTAGGTTGAGTTTAGACTTCTTGGTCGCGGCCAATGGCC |
| RW172 | ATATCCCGATTAACATAACATCCCACTTAGGAAGACTTTTT |
| RW249 | GTACCTGTTTCTTAACGACGATACAAGGAGGTAGTT |

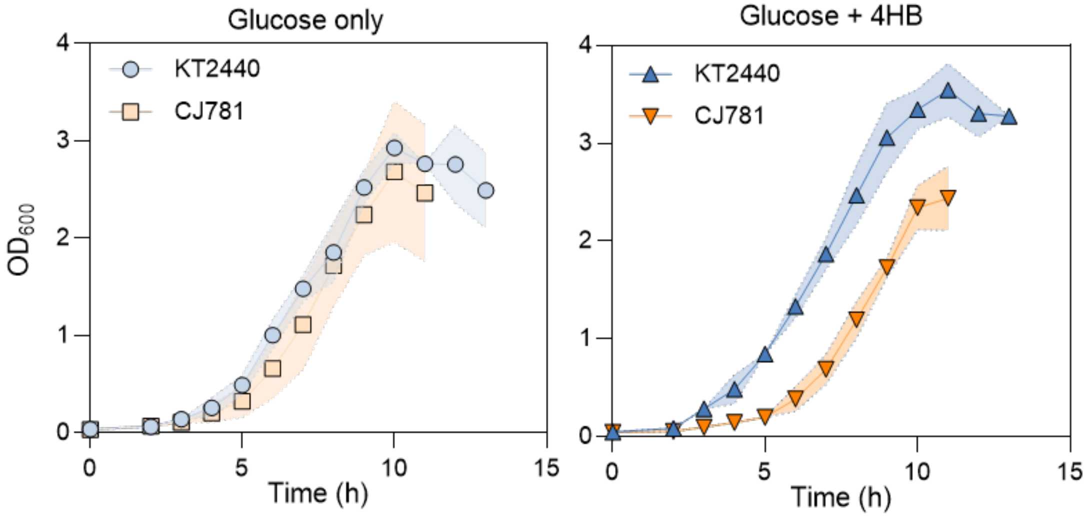

**Figure S1.** Growth dynamics of CJ781 and KT2440 on glucose alone or glucose plus 4HB. The absolute values are provided in **Excel file 1.** The data represent the mean ± the standard deviation determined from six independent biological replicates.

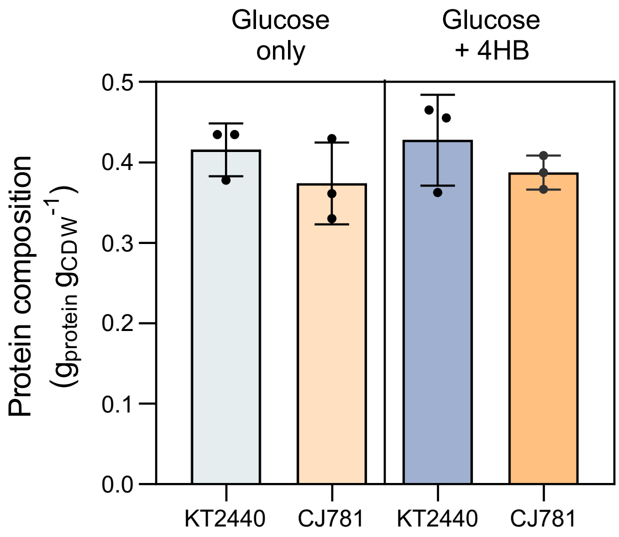

**Figure S2.** Protein content (g) per cell dry weight (g). Protein was measured using a bicinchoninic acid (BCA) protein assay. The data represent the mean ± the standard deviation determined from three independent biological replicates.

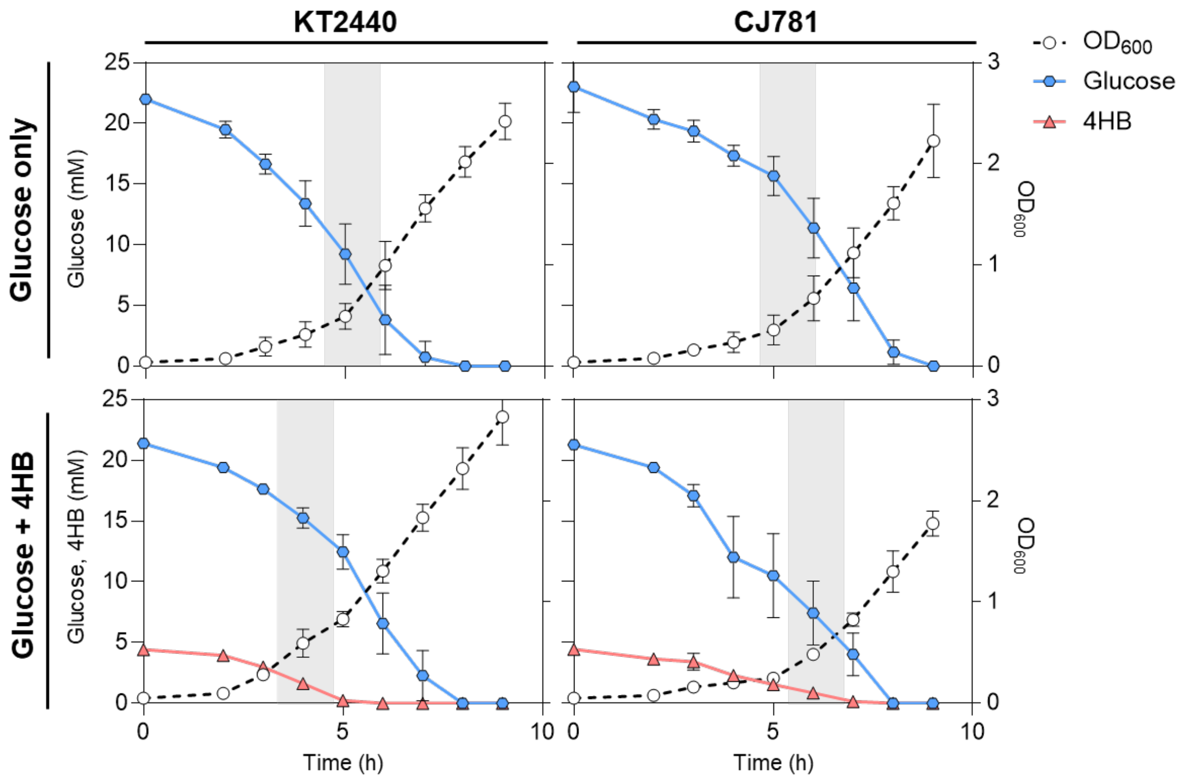

**Figure S3.** Kinetic profiles of substrate depletion and cellular growth ($OD_{600}$**).** The data represent the mean ± the standard deviation determined from three independent biological replicates. The shaded area represents the $OD_{600}$ range that was sampled for metabolomics, proteomics, and $^{13}C$-fluxomics.

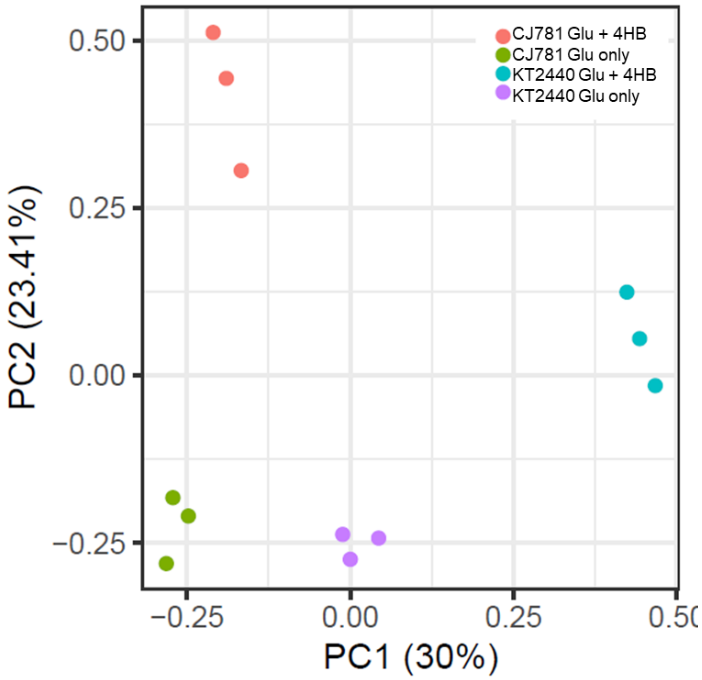

**Figure S4.** Principal component analysis of the strains and growth conditions.

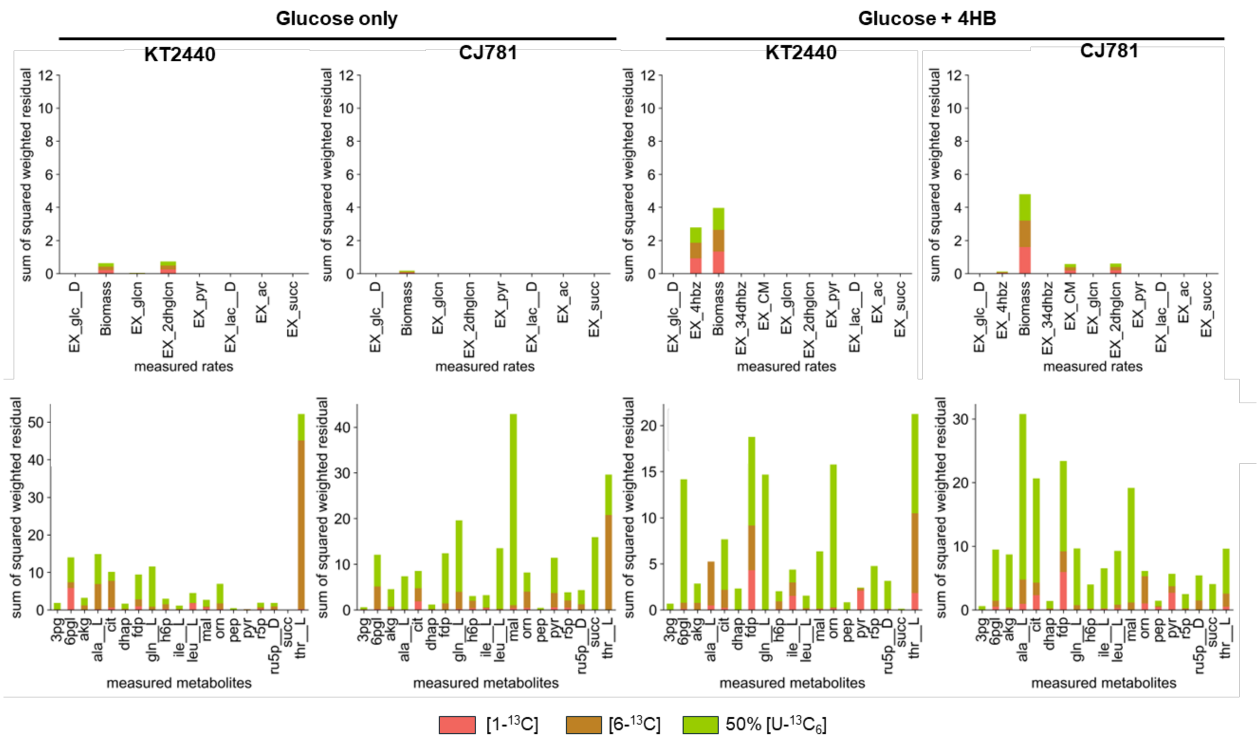

**Figure S5.** Sum of squared weighted residuals for each $^{13}$C-metabolic flux model. The model was optimized across parallel labeling experiments using three $^{13}$C-glucose substrates (100% [1-$^{13}$C], 100% [6-$^{13}$C], and 50:50% [U-$^{13}C_6$] and unlabeled) and three independent biological replicates.

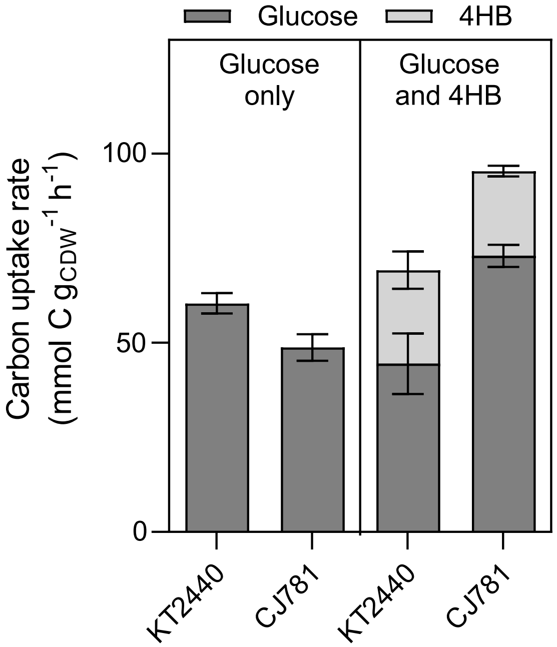

**Figure S6.** Uptake rates of glucose and 4HB, in mmol carbon $g_{CDW}^{-1}$ $h^{-1}$, calculated during exponential growth**.** The data represent the mean ± the standard deviation determined from three independent biological replicates.

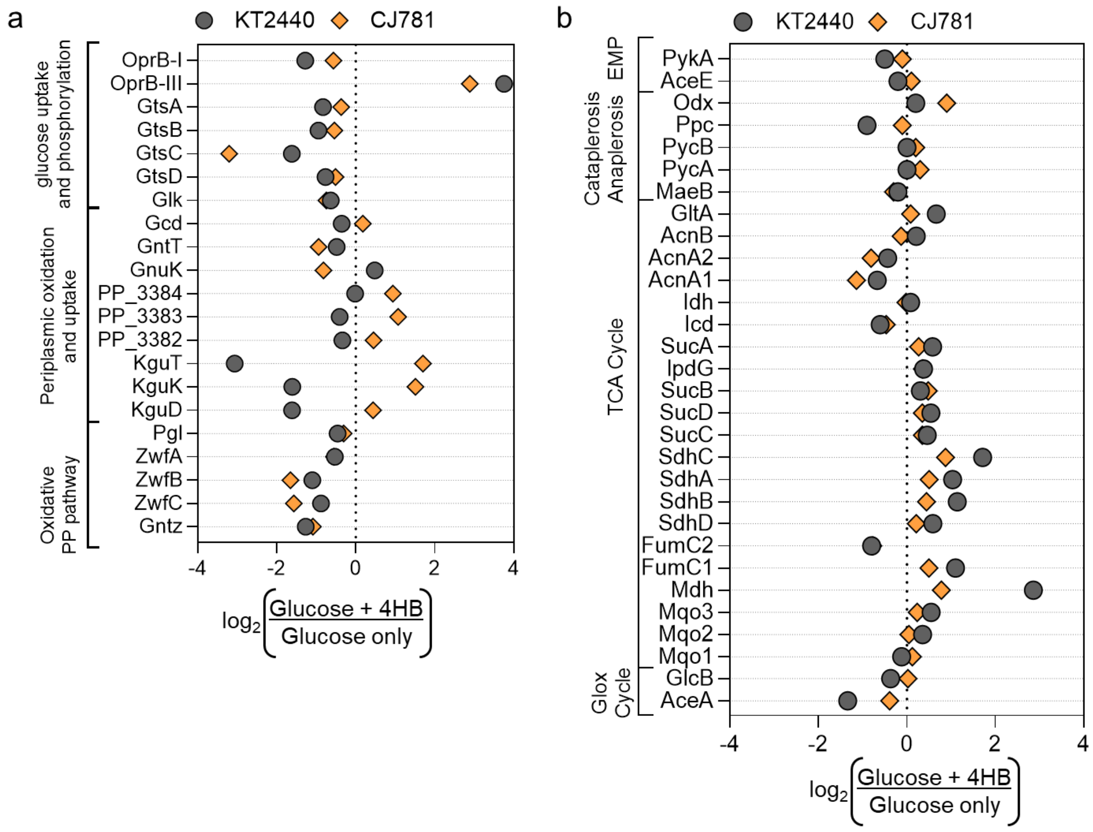

**Figure S7.** Differential protein abundance of central carbon metabolism enzymes for strains fed glucose plus 4HB relative to glucose alone. Proteins were measured from three independent biological replicates. Data is provided in **Excel file 1.**

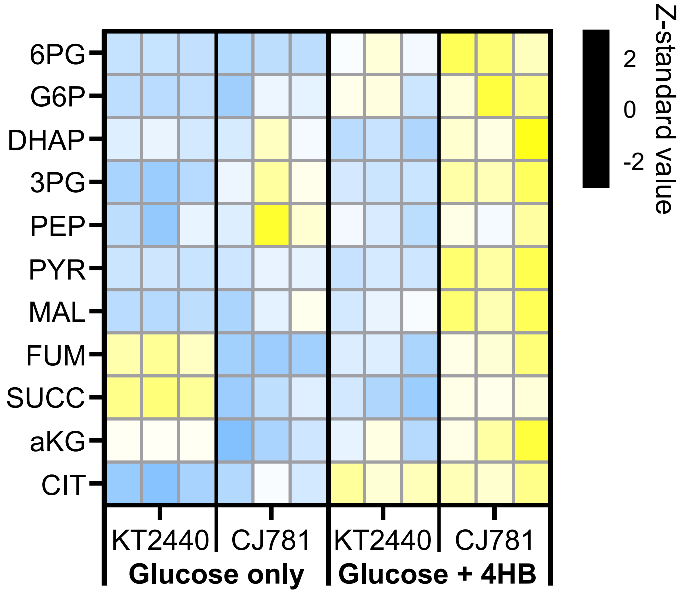

**Figure S8.** Heatmap of measured intracellular metabolites. The three independent biological replicates of each strain and growth condition were corrected by corresponding $g_{CDW}$ and normalized using Z-standard value to linear transform the measurements for each metabolite.

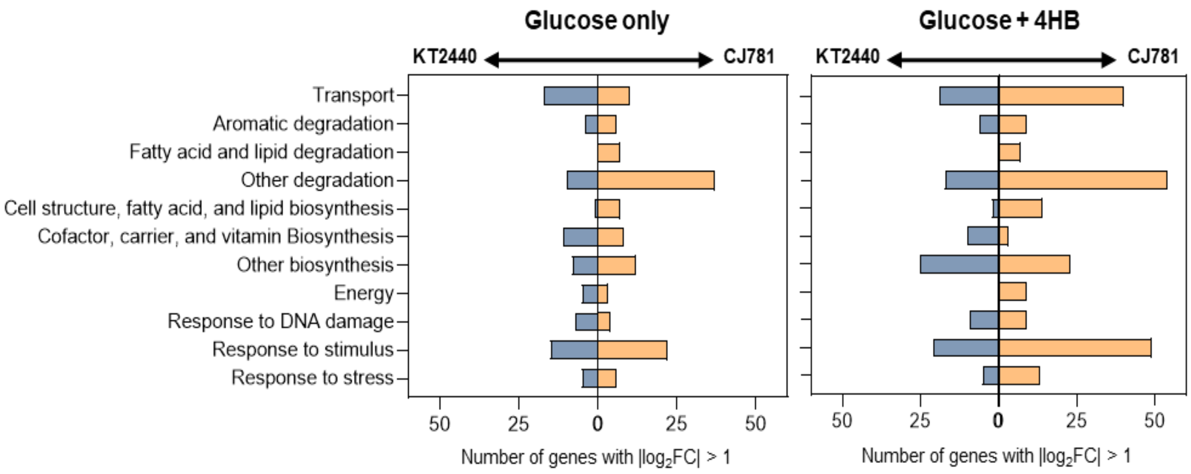

**Figure S9.** Global changes to the proteome in response to engineering muconate production. The data represent three independent biological replicates.

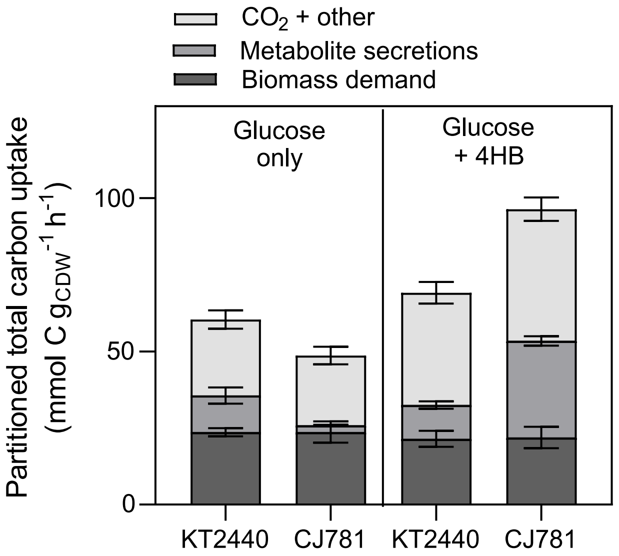

**Figure S10.** Metabolic investment of total substrate uptake, in carbon, into biomass generation, metabolite secretions, and $CO_2$. The data represent the mean ± the standard deviation determined from three independent biological replicates.

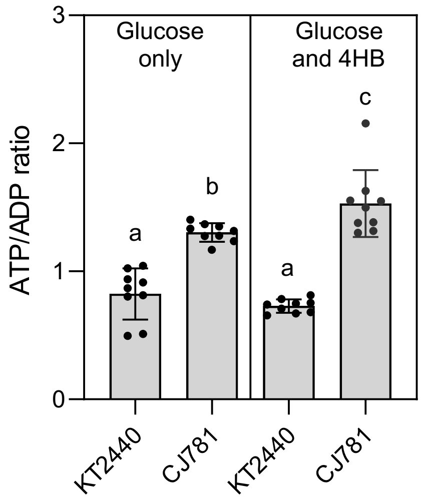

**Figure S11.** ATP to ADP ratio for the two strains and growth conditions. Error bars represent the standard deviation of the mean across three independent biological replicates and three technical replicates.

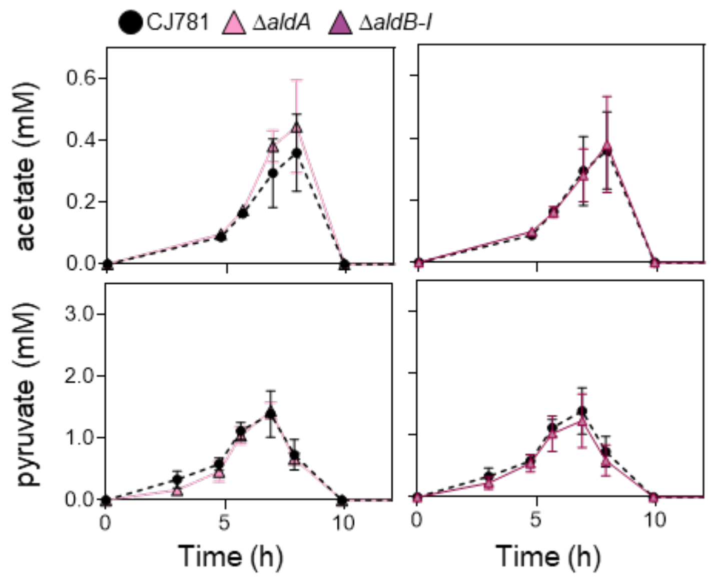

**Figure S12.** Impact of individual deletions of *aldB-I* and *aldA* in CJ781 on acetate and pyruvate secretions. The data represent the mean ± the standard deviation determined from three independent biological replicates.

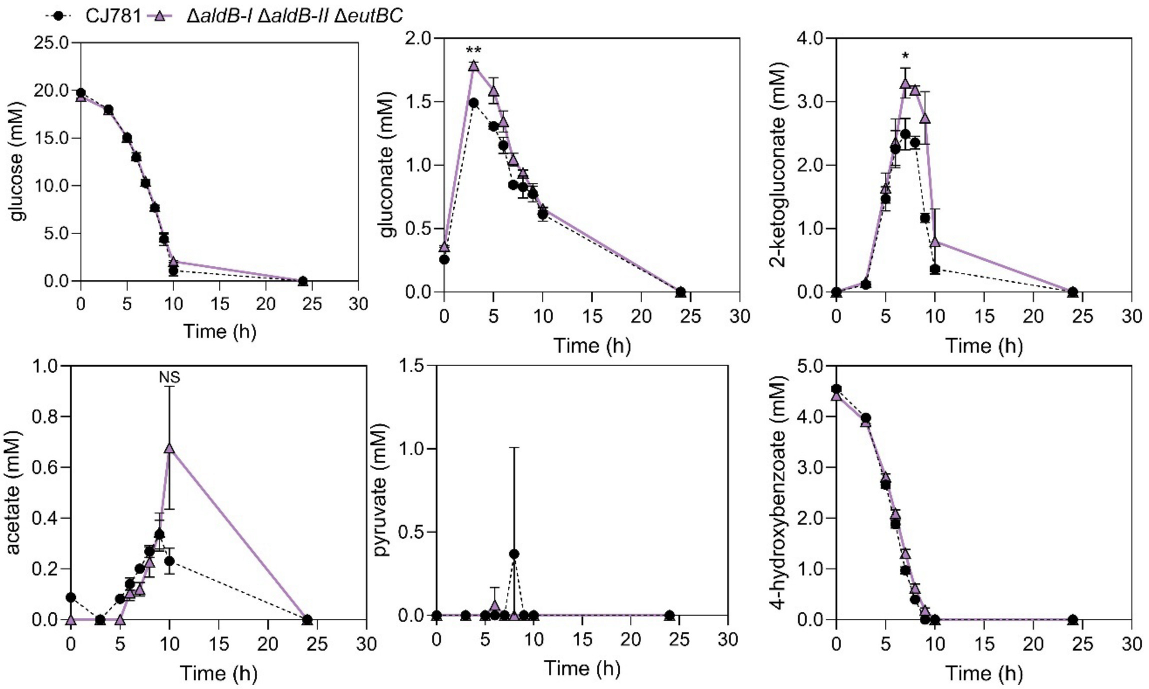

**Figure S13.** Impact of triple deletion of *aldB-I, aldB-II,* and *eutBC* in CJ781 on secreted metabolites. The data represent the mean ± the standard deviation determined from three independent biological replicates. Strain genotypes are provided in **Table 1.**

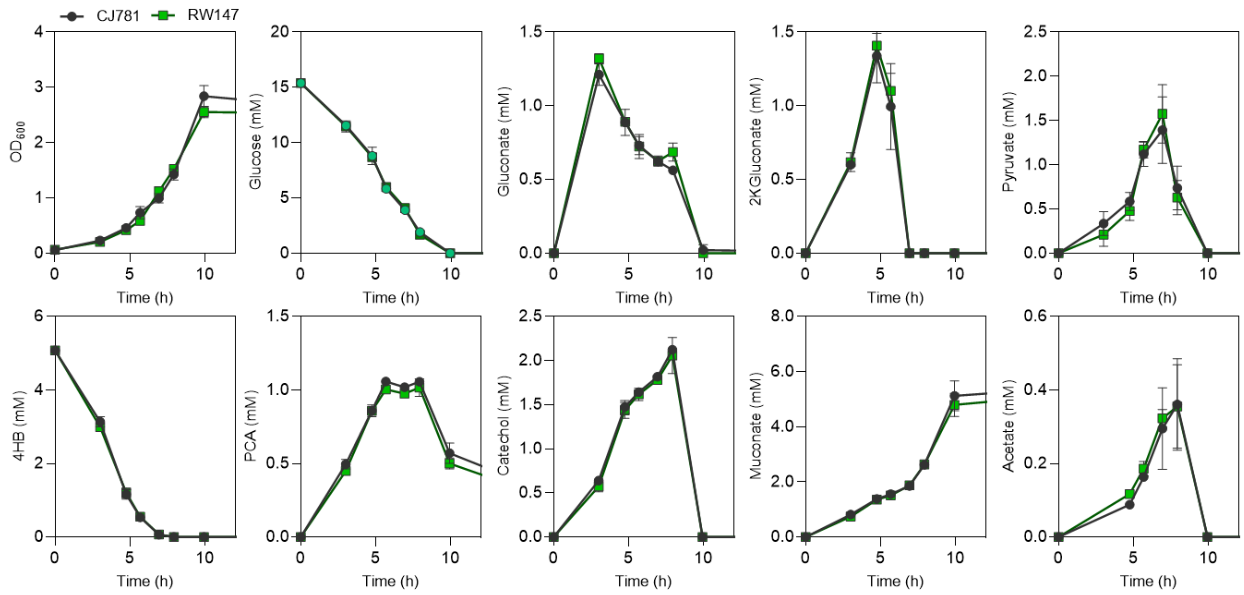

**Figure S14.** Comparison of CJ781 to RW147 (CJ781 with SAGE integration sites). The data represent the mean ± the standard deviation determined from three independent biological replicates.

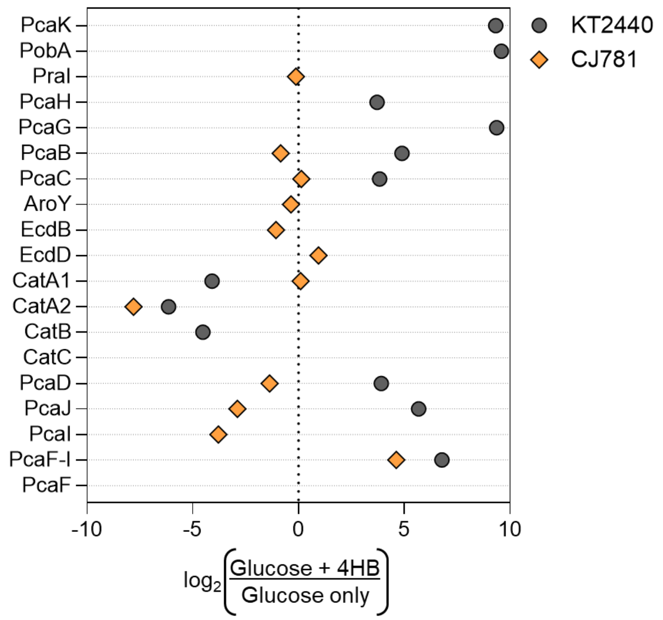

**Figure S15.** Differential protein abundance of aromatic pathway enzymes for strains fed glucose plus 4HB relative to glucose alone. Proteins were measured from three independent biological replicates. Data is provided in **Excel file 1.**

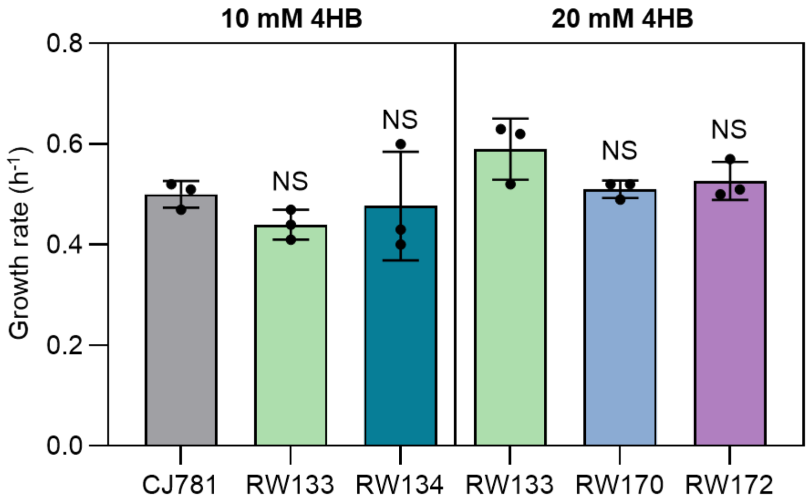

**Figure S16.** Growth rates of the aromatic pathway engineered strains fed 20 mM glucose plus either 10 mM 4HB or 20 mM 4HB. Statistically significant differences ($p$ < 0.05) were determined with an unpaired two-tailed $t$-test. The data represent the mean ± the standard deviation determined from three independent biological replicates.

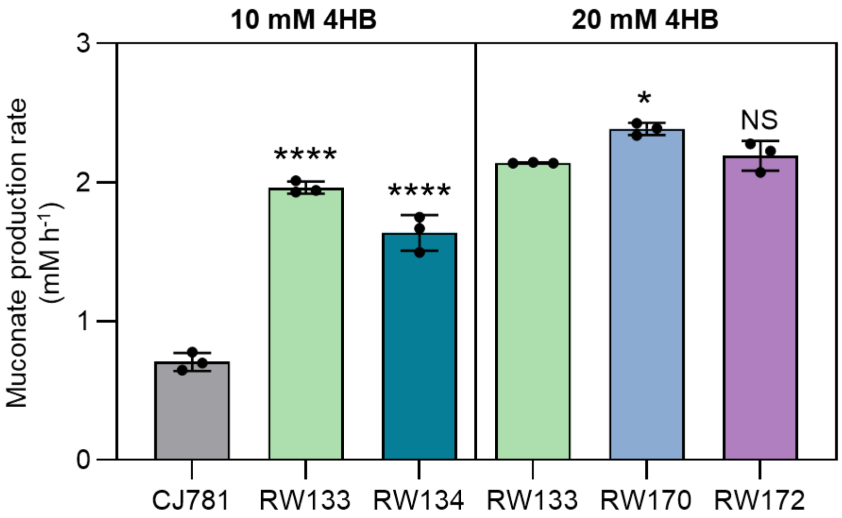

**Figure S17.** Muconate production rates quantified in shake flask experiments for the aromatic pathway engineered strains fed 20 mM glucose plus either 10 mM 4HB or 20 mM 4HB. Statistically significant differences ($p < 0.05$) were determined with an unpaired two-tailed *t*-test. The data represent the mean ± the standard deviation determined from three independent biological replicates.

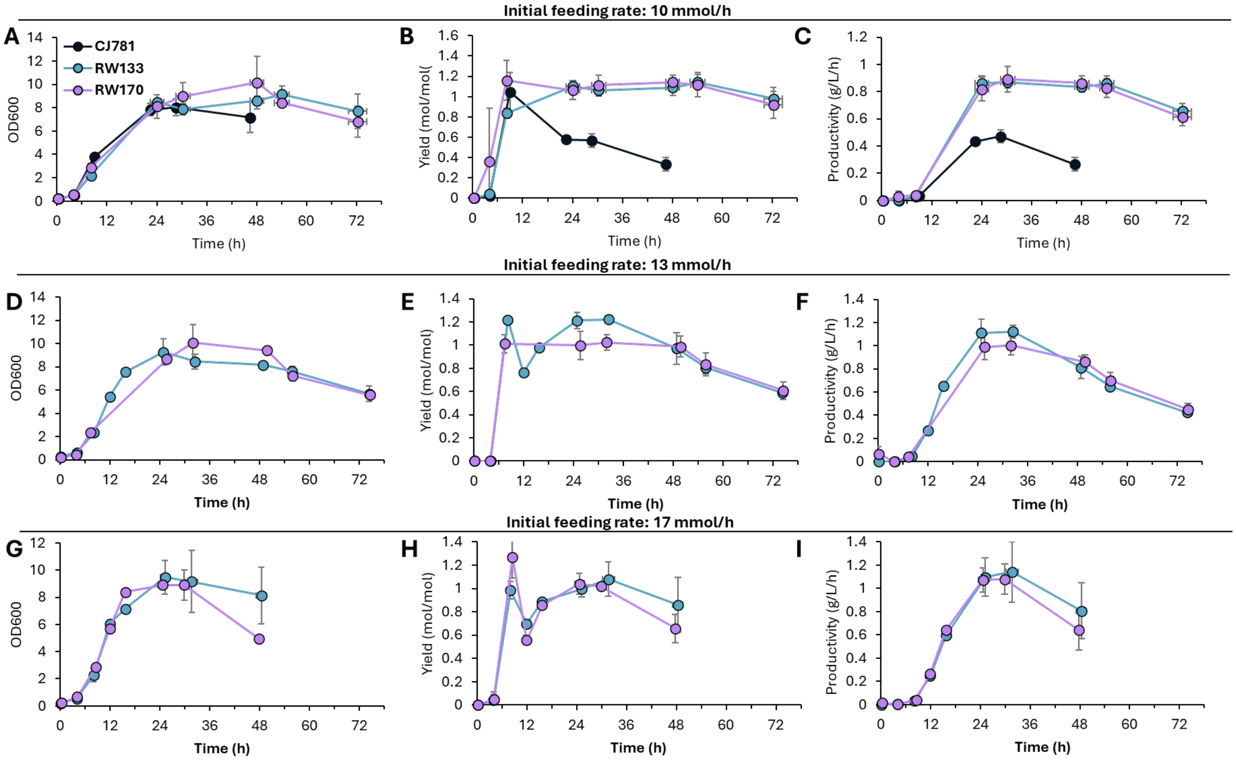

**Figure S18. Bacterial growth, muconate yields, and muconate productivities of CJ781, RW133, and RW170 in bioreactors at three different feed rates**. 4HBA feeding rates were maintained at (A,C) 10 mmol/h, (D–F) 13 mmol/h, and (G-I) 17 mmol/h, normalized to the batch volume. (A, D, G) Bacterial growth evaluated as OD600, (B, E, H) muconate yield (mol/mol) over time, and (C, F) muconate producivitity (g/L/h) over time by strains CJ781, RW133, and RW170. Data represent the mean of biological duplicates, except for RW170 at the 10 mmol/h feeding rate and bot RW133 and RW170 at the 17 mmol/feed rate, which were conducted in triplicate. Vertical error bars represent the absolute error between biological duplicates or the standard deviation among triplicates. Horizontal error bars represent the absolute time (h) difference between biological duplicates or the standard deviation among triplicates. Note: The data points corresponding to 12 h and 16 h in RW133 at 13 mmol/h were obtained from a single replicate. The data points corresponding to 12 h and 16 h in RW133 at 17 mmol/h were obtained from a single replicate, and those in RW170 from two replicates.

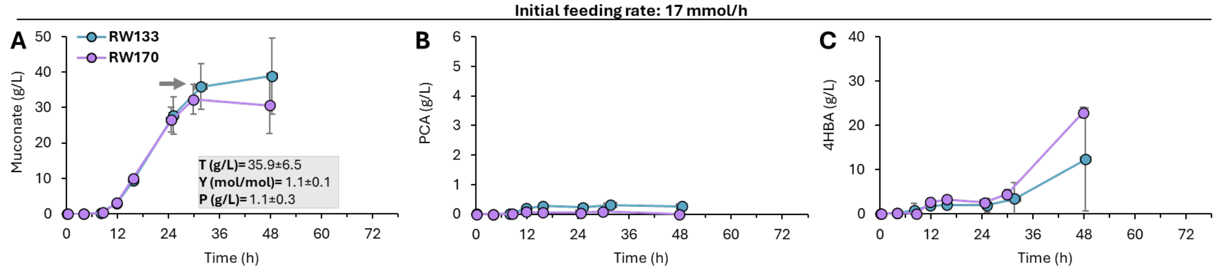

**Figure S19. Bioreactor profiles of CJ781, RW133, and RW170 at a feeding rate of 17 mmol/h.** 4HBA feeding rates were maintained at 17 mmol/h, normalized to the batch volume. (A) Muconate production, (B) protocatechuate (PCA) accumulation, and (C) 4HBA accumulation by strains RW133 and RW170. Grey arrow indicate the point at which titer (T), yield (Y), and productivity (P) were calculated before 4HBA accumulated substantially, with values shown in the grey box in the Figure. Data represent the mean of biological triplicates. Error bars represent the standard deviation among triplicates. Horizontal error bars represent the standard deviation of time (h) difference among triplicates. Note: The data points corresponding to 12 h and 16 h in RW133 were obtained from a single replicate, and those in RW170 from two replicates.
